# Fractional-order Approach to Modeling and Characterizing the Complex and Frequency-dependent Apparent Arterial Compliance: In Human and Animal Validation

**DOI:** 10.1101/2021.09.20.460769

**Authors:** Mohamed A. Bahloul, Yasser Aboelkassem, Taous-Meriem Laleg-Kirati

## Abstract

Recently, experimental and theoretical studies have revealed the potential of fractional calculus to represent viscoelastic blood vessel and arterial biomechanical properties. This paper presents five fractional-order models to describe the dynamic relationship between aortic blood pressure and volume, representing the apparent vascular compliance. The proposed model employs fractional-order capacitor element (FOC) to lump the complex and frequency dependence characteristics of arterial compliance. FOC combines both resistive and capacitive properties, which the fractional differentiation order, *α*, can control. The proposed representations have been compared with generalized integer-order models of arterial compliance. All structures have been validated using different aortic pressure and flow rate waveforms collected from various human and animal species such as pigs and dogs. The results demonstrate that the fractional-order scheme can reconstruct the overall dynamic of the complex and frequency-dependent apparent compliance dynamic and reduce the complexity. The physiological relevance of the proposed models’ parameters was assessed by evaluating the variance-based global sensitivity analysis. Moreover, the simplest fractional-order representation has been embed in a global arterial lumped parameter representation to develop a novel fractional-order modified arterial Windkessel. The introduced arterial model has been validated by applying real human and animal hemodynamic data and shows an accurate reconstruction of the proximal blood pressure. The novel proposed paradigm confers a potential to be adopted in clinical practice and basic cardiovascular mechanics research.

## I. INTRODUCTION

**C**ardiovascular diseases (CVDs) are the number one cause of premature death in the world. Decreased arterial compliance is recognized to be detrimental to the heart and arterial functions. Besides, variation in arterial compliance is associated with the major forms of CVDs, mainly hypertension and atherosclerosis [1]. Accordingly, assessment and evaluation of arterial compliance changes are crucial in diagnosing and treating hemodynamic disorders [2]. Vascular compliance stands for the ability of the vessel to store the blood. Functionally, it can be defined as the ratio of the incremental variation in the blood volume (*dV*) due to an incremental variation in distending pressure (*dP*). Accordingly, mathematically it is expressed as: *C* = *dV*/*dP* [3]. Over the last decades, several analytical and experimental studies have focused on modeling and characterizing vascular compliance [4], [5], [6], [7], [8], [9], [10], [11], [12], [13]. With the introduction of the well-known linear Windkessel representation of the arterial system, arterial compliance was assumed to have a single constant value for the entire cardiac cycle. Hence, the transfer function relating the blood volume variation to the blood pressure input changes was considered constant as well. Accordingly, the arterial compliance was modeled within the arterial lumped parameter circuit’s Windkessel as an ideal capacitor whose capacitance is constant [8]. However, this assumption was not realistic, and its drawbacks were reflected essentially in the estimation of the hemodynamic determinants [14]. In fact, it does not lead to the correct evaluation of the true value of arterial compliance [15]. Besides, by analyzing the transfer function blood volume/input pressure, experimental studies have shown that this relationship is frequency-dependent, and a time delay between the arterial blood volume and the input blood pressure is observed. Hence a variation in the arterial compliance along the cardiac cycle coexists [16], [3], [14].

In order to take into account this frequency dependence, some research investigations have promoted a new configuration where they considered the viscoelastic properties of the arterial vessel and represented the arterial compliance using the so-called *Voigt-cell* configuration[16], [17]. This type of arterial model was known as viscoelastic Windkessel. Although the viscoelastic *Voigt-cell* has resolved some contradictions of the standard elastic lumped parameter Wind-kessel, this configuration does also present some limitations as it does not account for the so-called stress-relaxation experiment [16]. To overcome this restriction, high-order viscoelastic configurations have been proposed by connecting many Voigt-cells. This solution might lead to an accurate estimation of arterial compliance and its feature; however, it is deemed a very complex alternative that poses extra challenges. Indeed, the number of parameters to estimate is more significant for higher-order models, while the obtained experimental data is habitually small and insufficient to identify all the parameters. It is also known that reduced-order models are more desirable for their uniformity and simplicity of investigation [18], [16].

In the last decades, non-integer differentiation, the so-called fractional-order differential calculus, became a popular tool for characterizing real-world physical systems and complex behaviors from various fields such as biology, control, electronics, and economics. The long-memory and spatial dependence phenomena inherent to the fractional-order systems present unique and attractive peculiarities that raise exciting opportunities to represent complex phenomena that represent power-law behavior accurately. Regarding cardiovascular modeling, the power-law behavior has been demonstrated in describing human soft tissues visco-elasticity and characterizing the elastic vascular arteries. The *in-vivo* and *in-vitro* experimental studies have pointed that fractional-order calculus-based approaches are more decent to precisely represent the hemodynamic; the viscoelasticity properties of soft collagenous tissues in the vascular bed; the aortic blood rate [19], [20]; red blood cell (RBC) membrane mechanical properties [21] and the heart valve cusp [22], [23], [24], [20].

In addition, recently, we developed novel fractional-order arterial Windkessel representations, [25], [26]. The proposed framework takes advantage of the relevant fractional-order calculus tools. Basically, the fractional-order Windkessel representations are similar to the well-known Windkessel configurations; however, instead of representing the arterial compliance with an ideal capacitor which is purely a storage element, we investigate the use of the fractional-order element, namely the fractional-order capacitor. Our elemental investigations in the frequency domain showed that the proposed models accurately reconstruct the arterial impedance and solve the hemodynamic inverse problem by estimating the different vascular biomechanics determinants. Moreover, a clear association between the central hemodynamic parameters and the fractional differentiation order (*α*) has been observed. In this context, fractional orders have been employed to describe and control the transition between viscosity and elasticity.

This paper presents five fractional-order model representations to describe the apparent vascular compliance by representing the active relationship between blood pressure and volume. Each configuration incorporates a fractional-order capacitor element (FOC) to lump the apparent arterial compliance’s complex and frequency dependence properties. FOC combines both resistive and capacitive attributes within a unified component, controlled through the fractional differentiation order, *α*. Besides, the equivalent capacitance of FOC is inherently frequency-dependent, compassing the complex properties using only a few numbers of parameters. In order to evaluate the physiological relevance of the developed representations’ parameters and calibrate the models, variance-based global sensitivity analysis was evaluated. The proposed representations have been compared with generalized integer-order models of arterial compliance. Both models have been employed and verified using different central pressure and flow rate waveforms secured from human and animal subjects such as pigs and dogs. In addition, the simplest fractional-order model’s structure of the proposed arterial compliance that consists of a single fractional-order capacitor has been integrated within a well-known arterial Windkessel configuration to represent proximal and distal compliances. Accordingly, we developed a new modified arterial Windkessel. This model has been applied and validated using real central hemodynamic waveforms.

## II. PRELIMINARIES

### A. Apparent Compliance

The apparent compliance, *C_app_*, refers to the arterial bed’s capacity to store blood dynamically. Functionally, it corresponds to the transfer function between the blood volume (*V*) and input blood pressure (*P_in_*). Here, we present its mathematical derivation. Based on the conservation mass, the arterial blood flow pumped from the heart to the vascular bed (*Q_in_*) which can be written as:

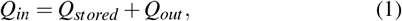

where *Q_stored_* is the blood stored in the arterial tree, and *Q_out_* corresponds to the flow out of the arterial system. In the frequency domain *Q_out_* can be expressed as:

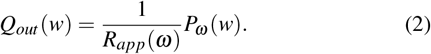

where *ω* corresponds to the angular frequency and *R_app_* is the apparent arterial resistance [3]. *Q_stored_* is defined as the rate of flow by taking the first derivative of the volume equation for the time.

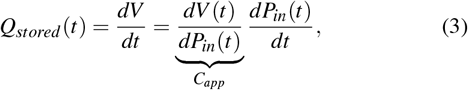

Hence in the frequency domain *Q_stored_* can be expressed as:

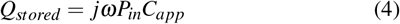

Aortic input impedance *Z_in_* defines the capacity of the vascular system to impede the blood rate dynamically. It corresponds to the left ventricular afterload. Functionally, it is expressed in the frequency domain, as the ratio between the arterial blood pressure (*P_in_*) and flow (*Q_in_*) at the aortic level of the systemic vascular system, that is:

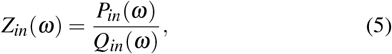

Substituting equations (2) and (4) into equation (1) gives:

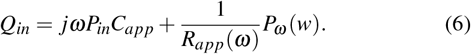

Rearranging the above equation yields an expression for *C_app_* in terms of *Zin* and *R_app_* as follow:

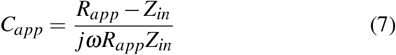

### B. Fractional-order capacitor

Fractional-order capacitor (FOC) known as the constant phase element [27] is a fractional-order electrical element representing the fractional-order derivative through its *curent-volatge* characteristic. In fact, the relationship between the current, *Q*(*t*), passing through an FOC and the voltage, *P*(*t*), across it with respect to time, *t*, can be written as follow:

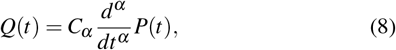

where *C_α_* is a proportionality constant so-called pseudocapacitance, expressed in units of [Farad/second^1−*α*^], [28]. The conventional capacitance, *C*, in unit of Farad is related to *C_α_* as *C* = *C_α_ω*^*α*−1^ that is frequency-dependent. The fractional-order impedance (*Z_α_*) is expressed as follow:

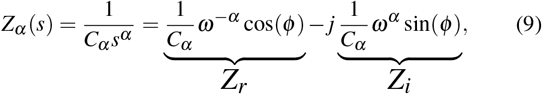

where s corresponds to the *Laplace* variable and *ϕ* denotes the phase shift expressed as: *ϕ* = *απ*/2 [rad] or *ϕ* = 90*α* [degree or °]. *Z_r_* and *Z_i_* are the real and imaginary parts of *Z_α_* corresponding to the resistive and capacitive portions, respectively. From (9), it is apparent that the transition between resistive and capacitive parts is ensured by *α*. If 0 ≤ *α* ≤ 1, the bounding conditions of *α* will corresponds to the discrete conventional elements: the resistor at *α* = 0 and the ideal capacitor at *α* = 1). As *α* goes to 0, (*Z_i_*) convergence to 0, and thus the fractional element looks like that a pure resistor, whereas as *α* goes to 1, (*Z_r_*) converges to 0 and hence, the fractional element serves as a pure capacitor, [29]. Fig. 1 (a) represents the schematic diagram for a FOC along with the ideal resistor and capacitor. Many studies have shown that FOC is equivalent to a resistor ladder network (RC tree circuit), [30], [31]. This structure is similar to the electrical analogy of the generalized Kelvin–Voigt viscoelastic model. Fig. 1 (b) presents the equivalent RC tree circuit of FOC of any order, and Fig. 1 (c) shows the equivalent RC tree circuit of FOC of order 0.5. Bearing these properties in mind, the fractional-order *α* parameter allows extra versatility in modeling viscoelastic systems [32].

**Fig. 1:**
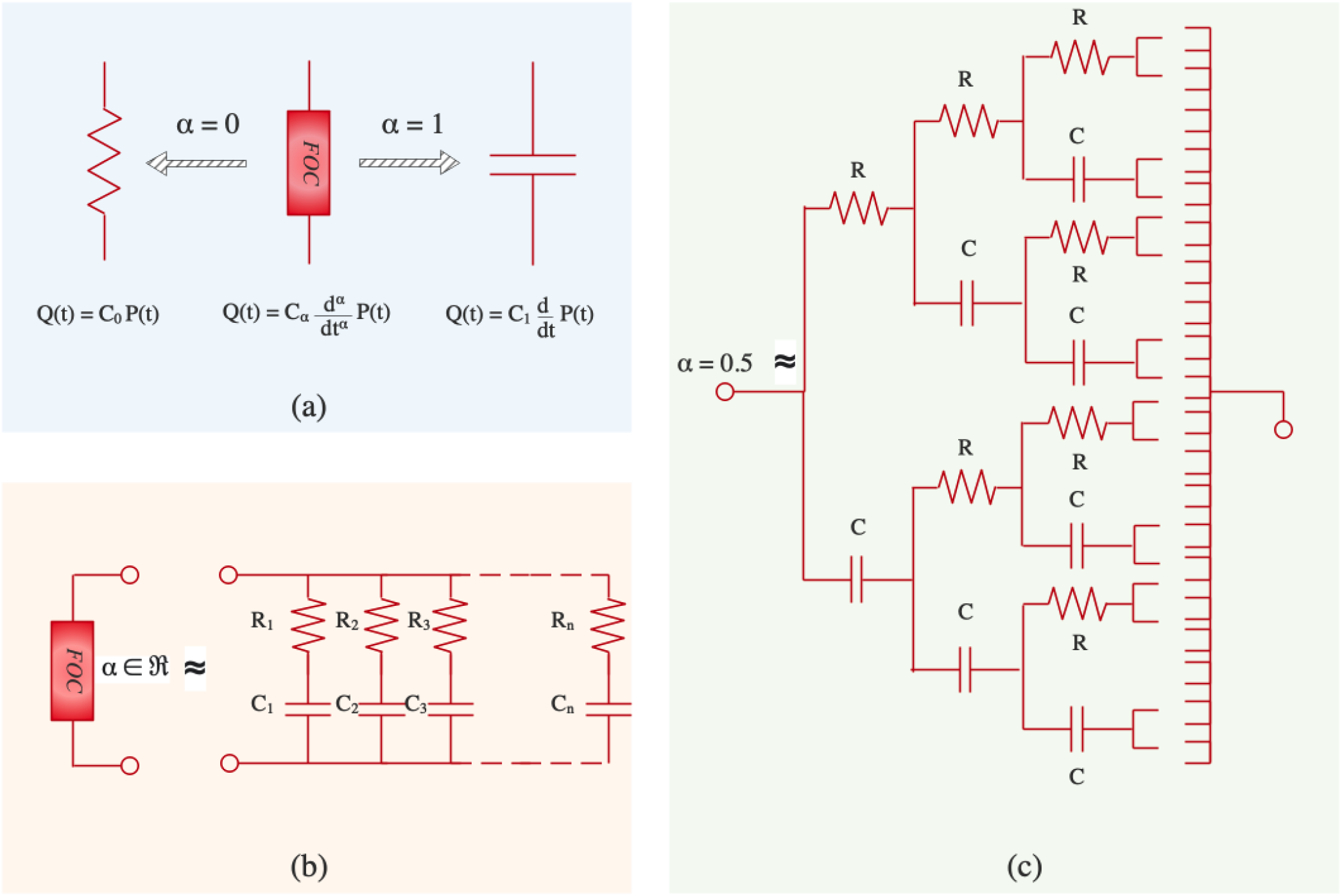
(a) Circuit diagram of the ordinary resistor and capacitor and the constant phase element is (fractional-order capacitor). It also shows the *current-voltage* relationship (*Q-P*): fractional-order capacitor; *Q*(*t*) = *C_α_*(*d^α^*/*dt^α^*)*P*(*t*) where 0 ≤ *α* ≤ 1 and *C_α_* is the pseudo-capacitance. The limits values of *α* namely for *α* = 0 and *α* = 1 corresponds to the ordinary elements the ideal resistor and capacitor respectively. (b) Circuit diagram representing the equivalent RC tree circuit of the fractional-order capacitor of any order, 0 < *α* < 1. (c) Circuit diagram representing the equivalent RC tree circuit of the fractional-order capacitor when *α* = 0.5.

## III. Models

As shown in the previous section, the FOC offers extra flexibility via its fractional differentiation order *α*, and it permits the smooth transition and control between the resistive and capacitive parts, which might be investigated to model the arterial system properties. By rewriting (3) in the fractional-order domain as:

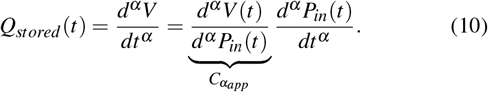

The FOC can be an inherent lumped element that can catch vascular compliance’s complex and frequency-dependent behavior. In fact, as expressed in (10), the pseudo compliance, *Cα_app_*, should be expressed in the unit of [1/mmHg .sec^1−*α*^] that makes, naturally, the standard compliance (*C_C_*), in the unit of [1/mmHg], frequency-dependent as:

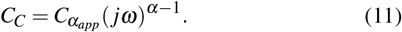

Hence the fractional-order capacitor presents physical bases in portraying the complex and frequency dependency of the apparent vascular compliance. Besides, based on the variation of the fractional differentiation order *α*, the real and imaginary parts of the resultant FOC’s impedance can possess various levels, so by analogy, *α* can control dissipative and storage mechanisms and hence the viscous and elastic component of the arterial wall. Furthermore, it is worth remarking that the equivalent circuit representation of FOC can be seen as an infinity Voigt cells branches joined in parallel. Consequently, FOC simplifies the representation of the complex arterial network’s mechanical properties by employing only two parameters (*α* and *C_α_*). In the following, we present the five fractional-order representations of arterial compliance shown in Fig. 2. In addition, as in this study, the proposed models were compared with generalized ordinary (integer-order) vascular compliance models; we also present their expressions following the fractional-order ones.

**Fig. 2:**
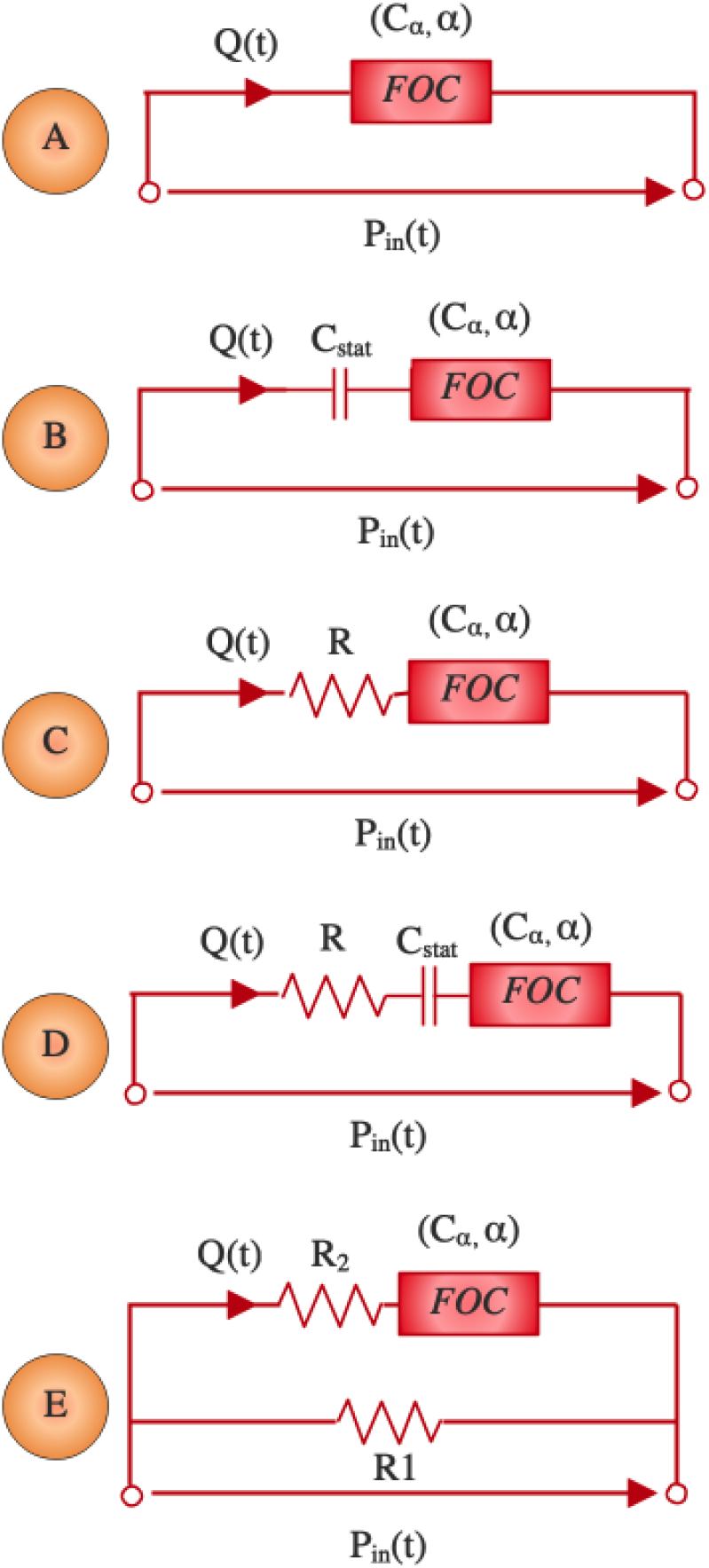
Schematic representations of the proposed fractional-order apparent compliance models.

### A. Fractional-order Models

#### Model A

As shown in Fig. 2, this representation consists of a single fractional-order capacitor. Accordingly, as mentioned previously, the apparent arterial compliance formulated in the unit of [1/mmHg] can be expressed as follows:

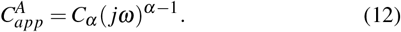

#### Model B

As shown in Fig. 2, this representation consists of an ideal integer-order capacitor (*C_stat_*) accounting for the static compliance connected in series to and *FOC*. The apparent arterial compliance formulated in the unit of [1/mmHg] can be expressed as follows:

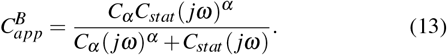

#### Model C

As shown in Fig. 2, this representation consists of an ideal resistor (R) connected in series to *FOC*. The apparent arterial compliance formulated in the unit of [1/mmHg] can be expressed as follows:

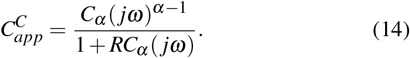

#### Model D

As shown in Fig. 2, this representation consists of an ideal resistor (*R*), an ideal integer-order capacitor (*C_stat_*), and FOC all connected in series. The apparent arterial compliance formulated in the unit of [1/mmHg] can be expressed as follows:

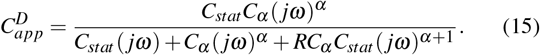

#### Model E

As shown in Fig. 2, this representation consists of an ideal resistor (*R*_1_) connected in parallel to a branch of a FOC in series with an ideal resistor (*R*_2_). The apparent arterial compliance formulated in the unit of [1/mmHg] can be expressed as follows:

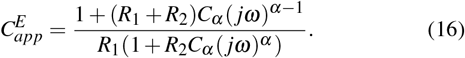

### B. Integer-order Models

#### Model F

This model expresses the apparent compliance based on the general viscoelastic model [35]. It is formulated as follows:

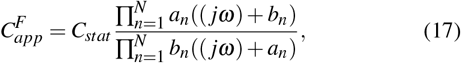

where *a_n_* and *b_n_* corresponds to imperial constants that can be adapted to fit any special case. *C_stat_* expresses the static compliance. Goedhard et. all pointed that this model could fit real experimental data with *N*=4, which we adopt in our comparative study.

#### Model G

It corresponds to the Voigt-cell based-representation. It consists of an integer-order, ideal capacitor (*C_stat_*) accounting for the static compliance in series to a resistor (*R_d_*), accounting for viscous losses.

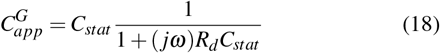

## IV. Method & Material

### A. In-vivo human and animal datasets

In order to validate the proposed approach, in this study, we use real data for human aging human hypertension and animal subjects. The *in-vivo* human data was extracted and digitized from aging, and hypertensive studies (Nichols et al., [33]). The aging human data consists of measured aortic blood flow rate (*Q_a_*) and aortic blood pressure (*P_a_*) at various ages, specifically 28, 52, and 68 years whose cardiac cycle is *T* = 0.95 *Sec*. The hypertensive pulse waves correspond tc the aortic blood flow rate and pressure waveforms for three human subjects suffering from three different hypertension stages, particularly normotensive, mild-hypertension, and severe hypertension. Their cardiac cycle is *T* = 0.92 *Sec*. In addition, to further compare and check the efficiency of the proposed models, we utilized animal (pigs and dogs) data extracted and digitized from (Segers et al.,[34]).The cardiac cycle of the dog is *T* = 0.46 Sec, and the pig’s one is *T* = 0.52 *Sec*. Figures 3–5 show the blood flow and pressure signals for each subject that we have used.

**Fig. 3:**
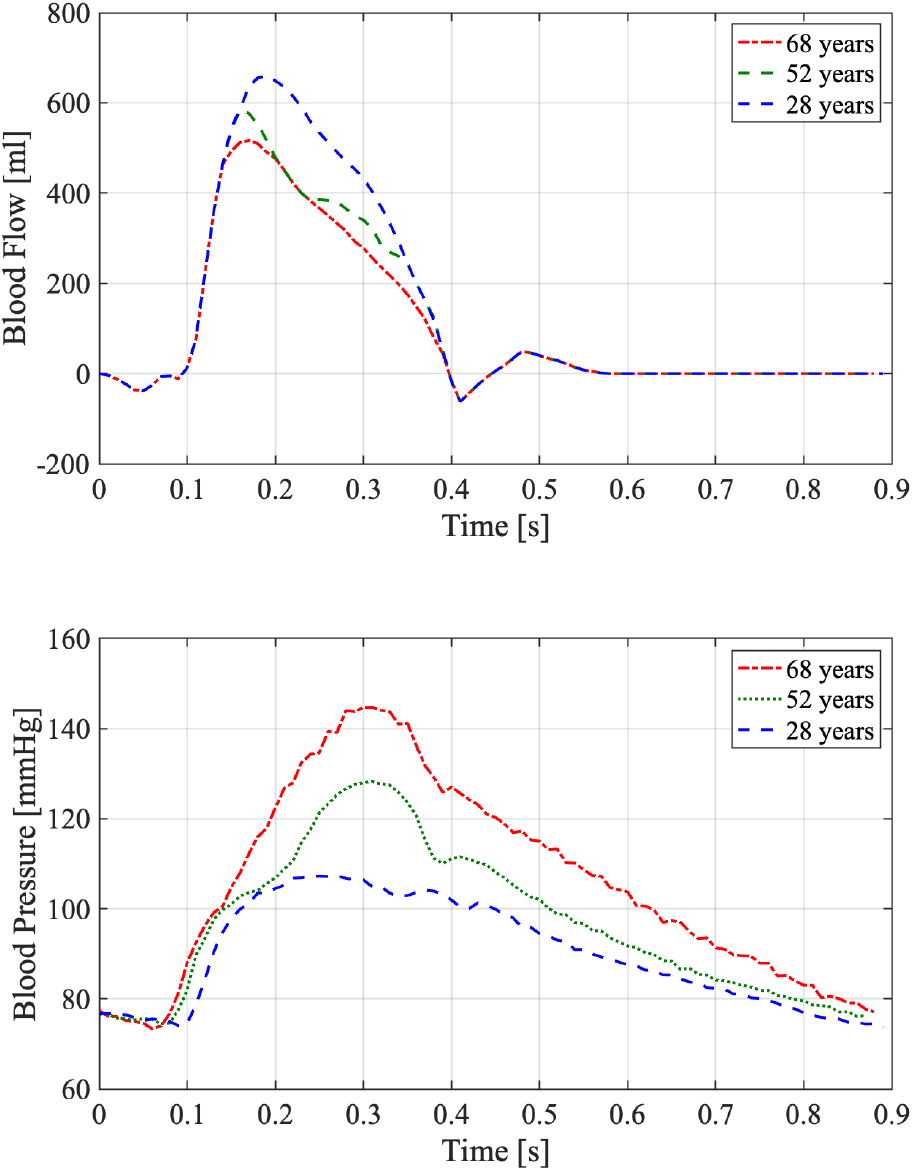
*In-vivo* human aging data sets digitized from ( Nichols et al., [33]). It presents the blood flow rate and pressure waveforms at different ages, namely 28, 52, and 68 years.

**Fig. 4:**
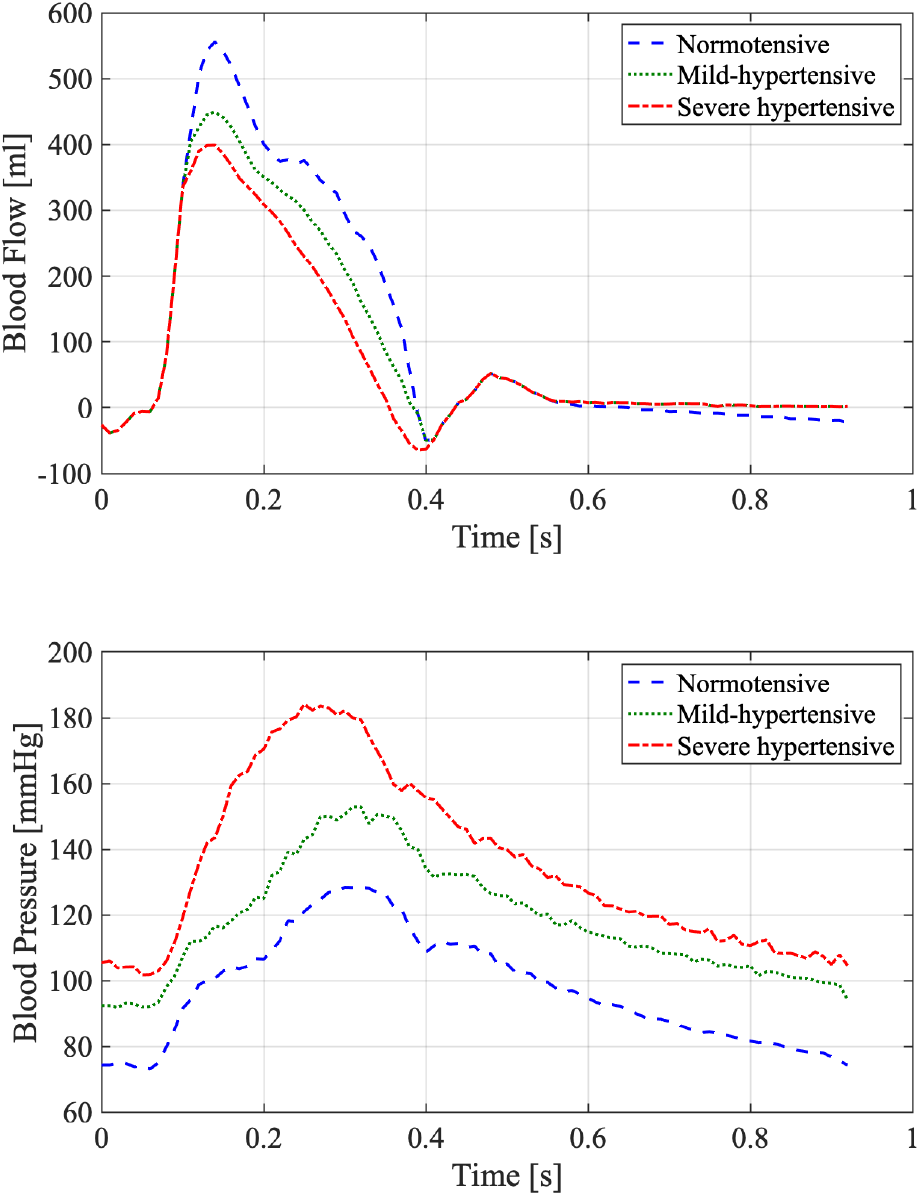
*In-vivo* human hypertension data sets digitized from (Nichols et al., [33]). It presents the blood flow rate and pressure waveforms at different hypertension conditions, namely normotensive, mild-hypertension, and severe-hypertension.

**Fig. 5:**
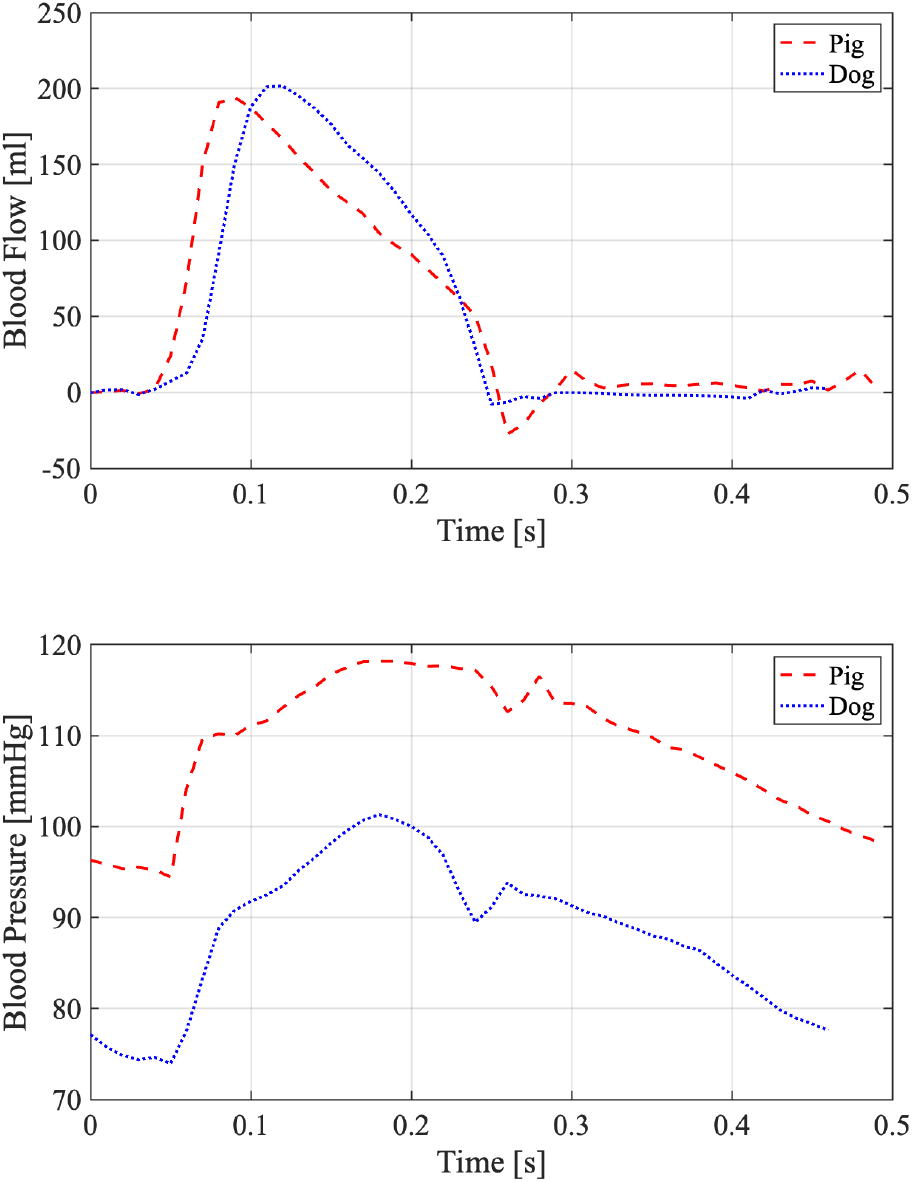
*In-vivo* animal (Pigs and Dogs) data sets digitized from (Segers et al.,[34]).

### B. Sensitivity analysis for the proposed arterial apparent compliance

In order to study how the variation in the apparent arterial compliance modulus and phase is associated with the variations of the different input parameters factors, a global sensitivity analysis based on *variance method* has been performed. Variance-based Sensitivity Analysis (VBSA) is a valuable step in the model calibration process, estimating the model parameters. In fact, it provides a relevant insight on how changes in the estimates of the parameters (the inputs of the model) map into variations of the performance metric that evaluates the model fit. A detailed review with practical workflow about the sensitivity analysis literature can be found in [36], [37], [38]. In this study, we evaluate the VBSA of the fractional-order arterial compliance using *First-order indices* known also as ‘*main effect*’ and the *total-order indices* so called ‘*total effect*’. The ‘*main effect*’ indices measure the direct contribution of the output variation from individual input factor or, equivalently, the expected reduction in output variance that can be obtained when fixing a specific input [36]. The *First-order indices* is defined as:

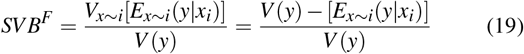

where *E* denotes expected value, *V* denotes the variance, x denotes the input, y denotes the output and *x* ~ *i* denotes “all input factors but the i-th”. The *total-order indices* evaluates the overall contribution from an input factor considering its direct effect and its interactions with all the other factors, which might amplify the individual effects. It is defined as:

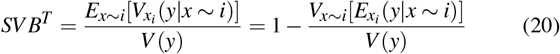

### C. Parameters fitting of the models

To fully identify the proposed fractional-order model and the integer-order based apparent compliance representations, the parameters and the fractional differentiation orders have to be estimated using the measured flow and pressure waveforms. The estimation process was based on a non-linear least square minimization routine applying the well-known MATLAB – R2020b, function *fmincon*. The parameters to estimate for each model’s representation 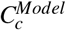 are refereed as *Θ^Model^* where *Model* = {(*A*), (*B*), (*C*), (*D*), (*E*), (*F*), (*G*)} denotes to the index of the model’s structure.

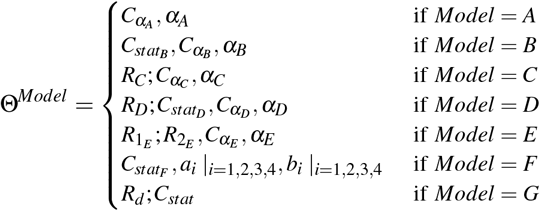

Fig. 6 exposes a flowchart describing the process of the models’ parameters estimation and performance analysis.

**Fig. 6:**
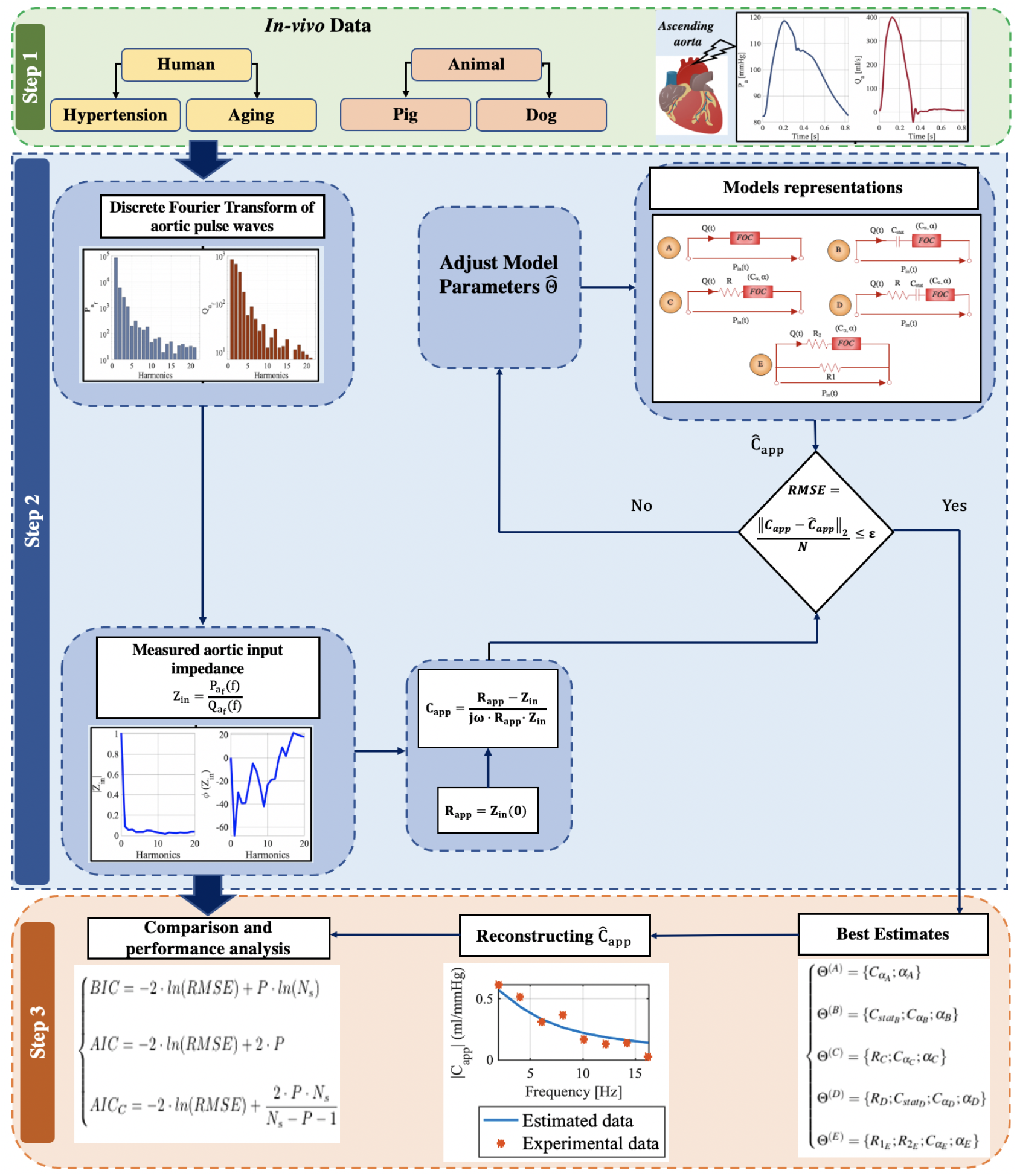
A flowchart showing the three main steps of the models’ calibration and performance analysis.

#### Algorithm 1 Parameter calibration of the models

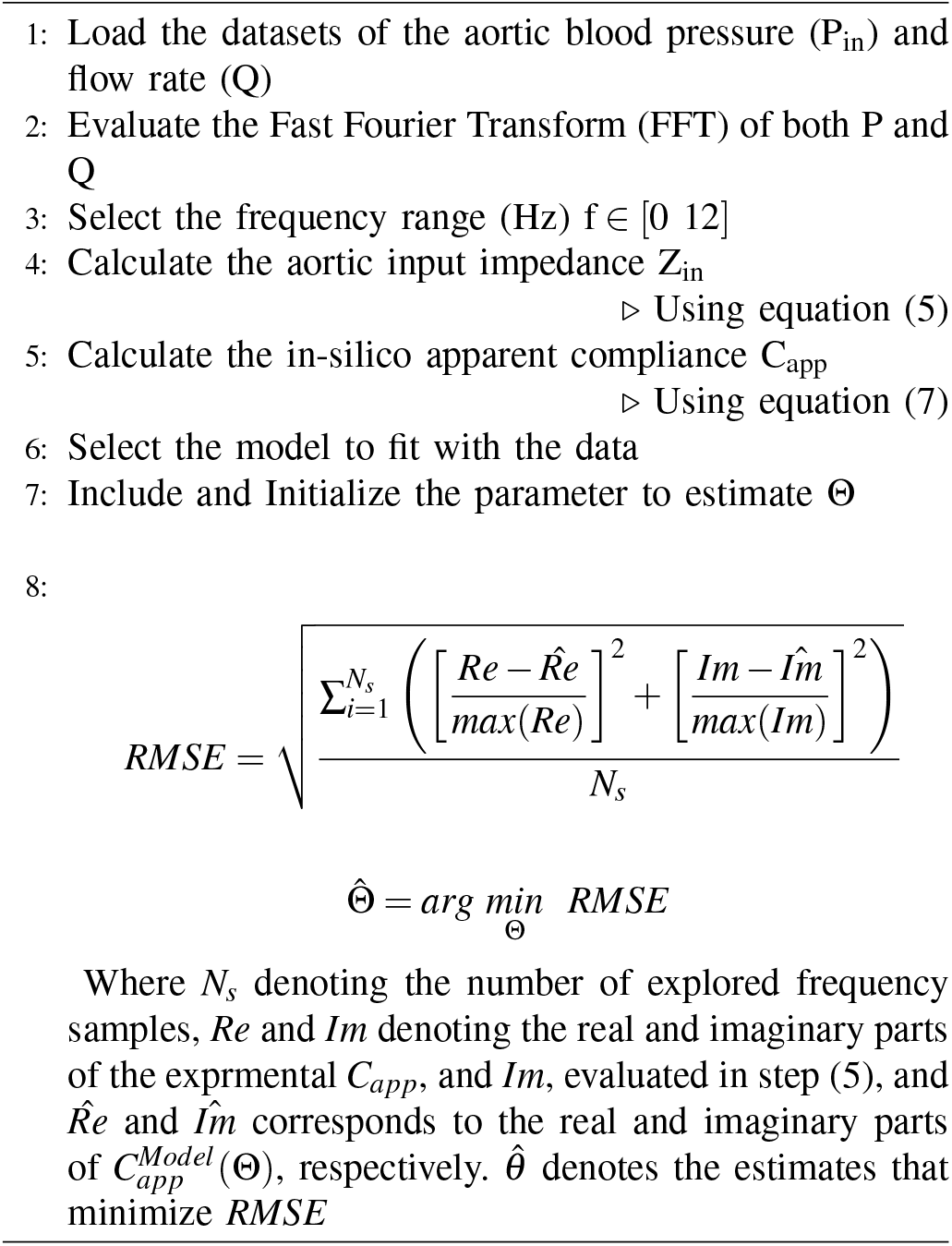

Furthermore, algorithm 1 summarizes the different steps applied to identify the different representations and estimate the model’s parameters.

### D. Statistical Analysis & convergence analysis

To analyze the ability of the developed models in reproducing the apparent arterial compliance dynamic, we evaluate the *RMSE*. In addition, because the model representations possess various numbers of parameters, to conduct a legitimate measurement and comparison, in addition to the *RMSE*, we assess the following criteria:

- *Bayesian Information Criterion, (BIC)*:

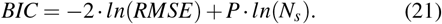
- *Akaike Information Criterion, (AIC)*:

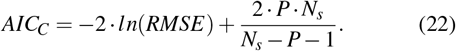
- *Corrected Akaike Information Criterion (AIC_C_)*:

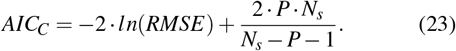

In the above expressions, *P* corresponds to the number of parameters.

Furthermore, to check the convergence of the estimates of the unknown parameters, we assess the time history of the estimated values of every fractional-order model during the optimization process starting from different initial conditions.

In addition, we show the time history of the objective function (RMSE) during the optimization process starting from different initial conditions.

## V. Results & Discussio

### A. Variance based sensitivity analysis in the fractional-order models

Fig. 7 shows the evaluated *VBSA*(*f*) based on main effect and total effect indices for the modulus of the complex and frequency-dependent apparent compliance for each fractional-order model. Fig. 8 displays the evaluated *VBSA*(*f*) based on main effect and total effect for the phase of the complex and frequency-dependent apparent compliance for each fractional-order model. In all cases, the *VBSA* was computed over a frequency range (Hz) *f* ∈ [0 30]. For the ease of visualization, for each model, all the parameters are listed in *y-axis*, whereas the *x-axis* represent the frequencies at which the main and total effect indices were computed. It is clear from these plots that the modulus and phase-based *Model (A)* and *Model (B)*are very sensitive to the fractional orders *α_A_* and *α_A_* respectively. With respect to the modulus, it worth to note that this sensitivity is minimal at low frequencies in favor of *C_α_A__* for *Model (A)* and (*C_statB_* = *C_α_B__*) for *Model (B)*. This result indicates that the initialization of the fractional differentiation orders for *Model (B)* and *Model (B)* should be set properly in the parameter’s calibration process. In addition, it marks that the fractional orders in these cases have central control in the variation of the modulus and phase of the apparent arterial compliance. Accordingly, this parameter might play an important function as a bio-marker assessing the transition between viscosity and elasticity; hence a potential index for arterial stiffness. With respect to *Model (C)* and *Model (D)*, generally the modulus and phase are not very sensitive to the variation of the fractional order differentiation *α_C_* and *α_D_*, respectively. The *α_C_*-related *VBSA* based on total effect is around 1 in the modulus case, and it increases with frequency concerning the phase. The *α_D_*-related *VBSA* based on total effect increases as the frequency increases for both modulus and phase. For *Model (E)*, it is noticeable that the fractional differentiation order *α_E_* plays an important role over the whole frequency range for both the phase and modulus of the apparent compliance. In addition, it is considered the most critical for the parameters estimation procedure. Conclusively, the sensitivity analysis based on the variance method was very informative to understand the role of the fractional-order over the frequency domain and to evaluate the effect of the parameters variation on the changes of the modulus and phase of the apparent compliance. This analysis is also very useful for the subsequent step consisting of the model calibration and real fitting data.

**Fig. 7:**
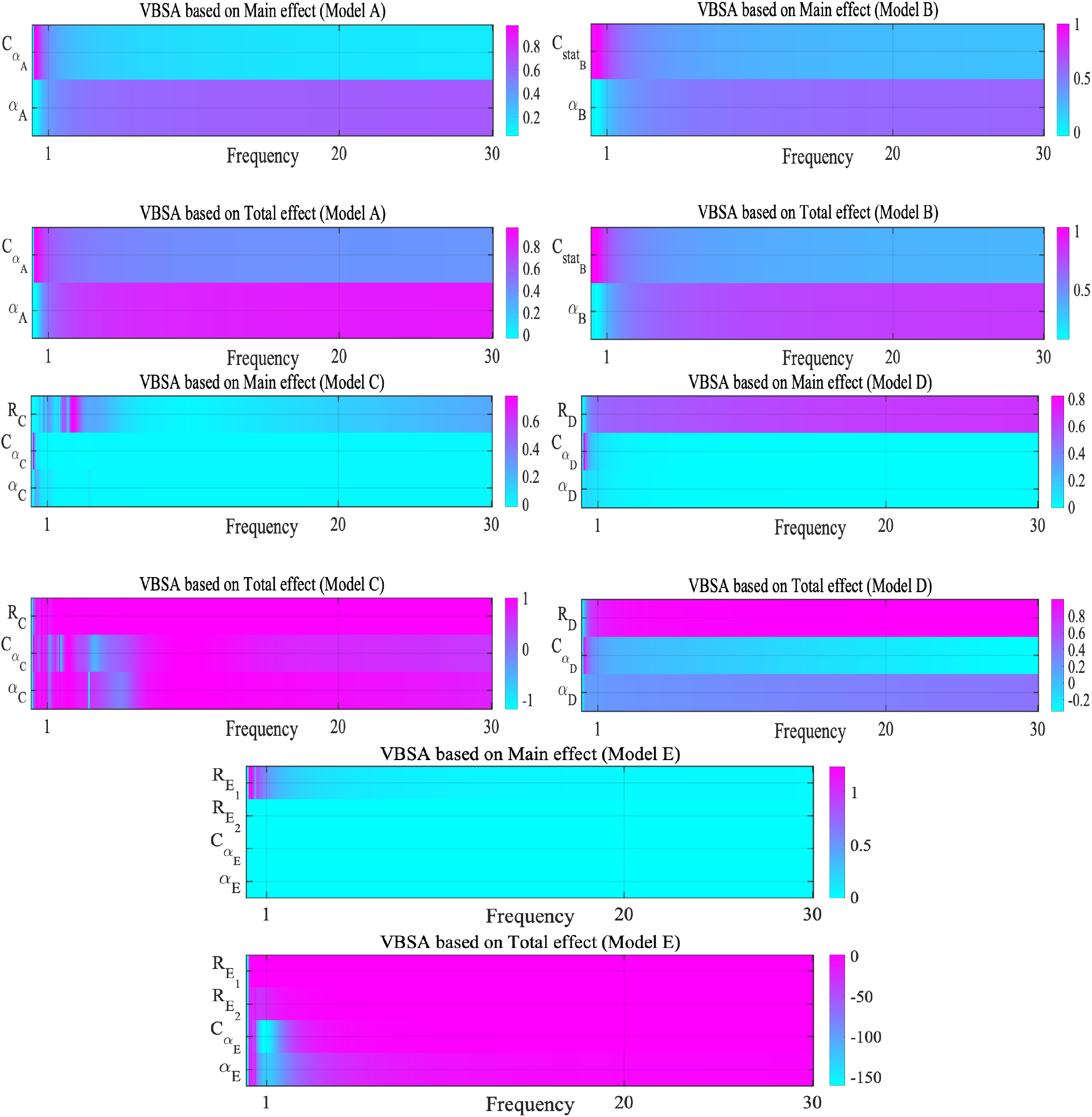
Main effect and total effect related-variance based sensitivity analysis with respect to the modulus of the fractional-order complex and frequency dependent apparent compliance

**Fig. 8:**
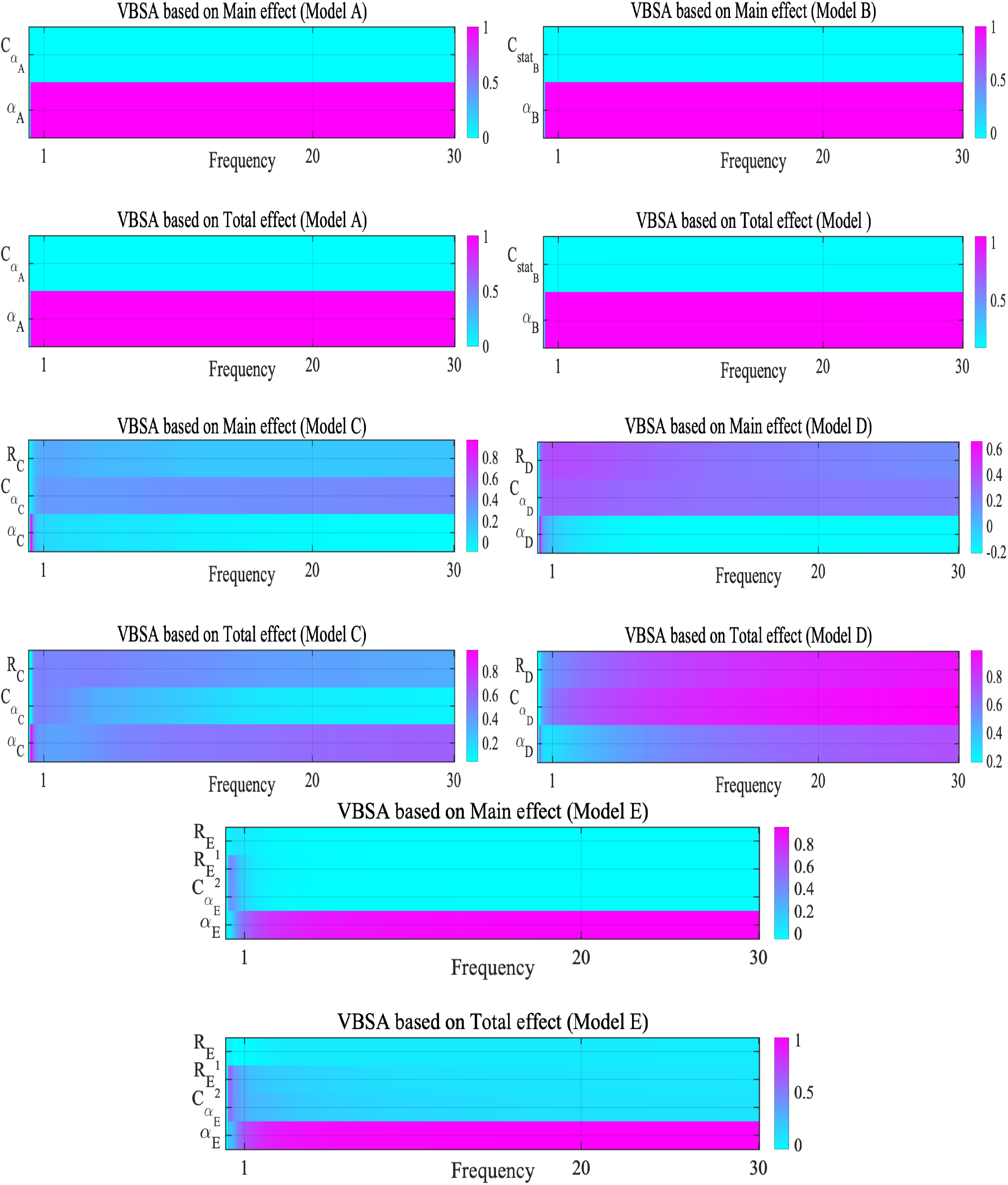
Main effect and total effect related-variance based sensitivity analysis with respect to the phase of the fractional-order complex and frequency dependent apparent compliance

### B. Model calibration

The evaluated *RMSE* and {*BIC*, *BIC* and *AIC_c_*}, values after applying the proposed model and integer-order ones to the experimental human and animal data, are presented in TABLE I–IV, respectively. The reconstructed apparent arterial compliance magnitudes using the fractional-order and integer-order models and the experimental magnitude are shown in Fig. 9. Each row in this figure corresponds to a specific subject from the real data, and each column corresponds to a particular model. Analyzing these results makes it clear that the fractional-order model representations grant an acceptable reproducing of the real arterial apparent compliance with a minimum number of parameters. The fractional-order *Model (C)* and *Model (E)* present the best RMSE comparing to the other fractional-order representation. The comparison between the proposed fractional-order models and the integer ones, namely Model (*F*) and (*G*) confirms that as the models differ in terms of performance, there is a trade-off between complexity induced by the number of parameters per representation and accuracy. Indeed, high-order models deliver high precision, however, at the expense of complexity. In order to take into account this compromise, the *BIC* and *AIC* criteria have been evaluated. Accordingly, *Model (A)* and *Model (B)* present the minimum values. Overall, using the fractional-order element enhances the accuracy of arterial compliance and reduces the complexity. For example, approximately the same performances are obtained using both *Model (C)* and *Model (F)*. However, in the representation-based *Model (C)*, only three parameters have been used rather than nine parameters in *Model (F)*.

**Fig. 9:**
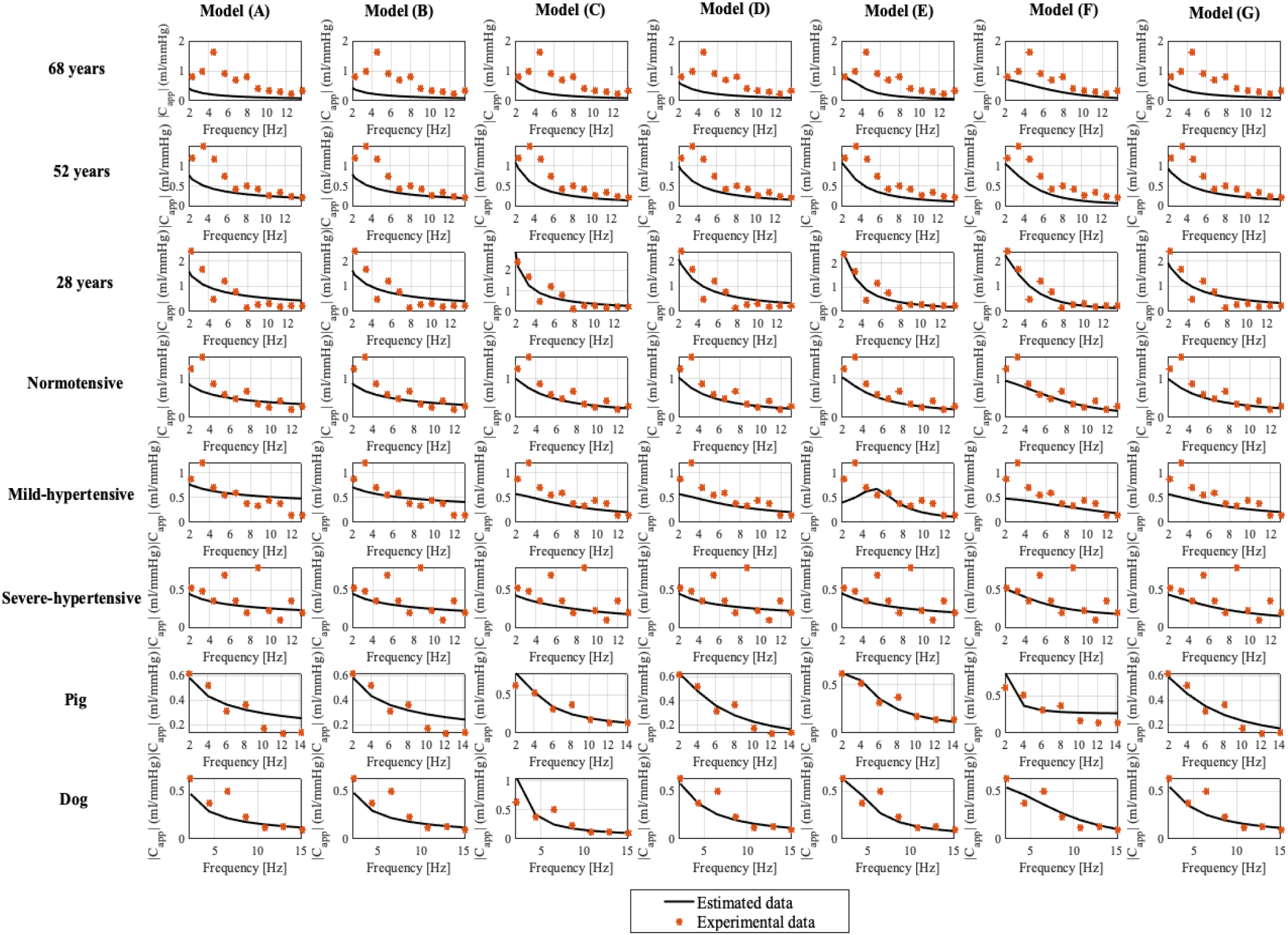
Estimated apparent compliance magnitudes using proposed fractional-order and standards apparent compliance models {Model (A), Model (B), Model (C), Model (D), Model (E), Model (F), Model (G)} along with the experimental *In-vivo* human-aging and animal (Pigs and Dogs) ones.

**TABLE I:**
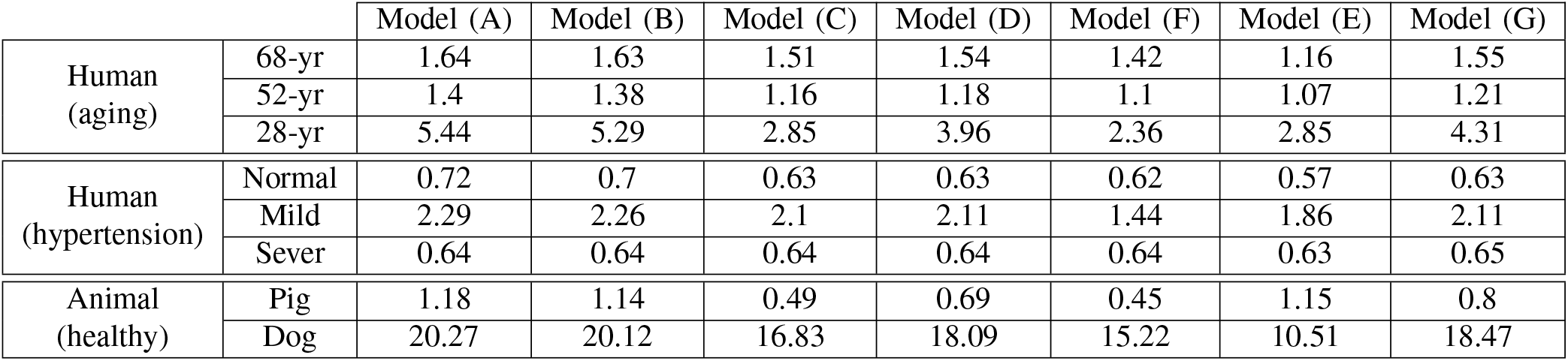
*RMSE* calculated based on the developed fractional-order representation and standard ones for each subject.

**TABLE II:** *BIC* calculated based on the developed fractional-order representation and standard ones for each subject.

|  |  | Model (A) | Model (B) | Model (C) | Model (D) | Model (F) | Model (E) | Model (G) |
| --- | --- | --- | --- | --- | --- | --- | --- | --- |
| Human (aging) | 68-yr | 4.14 | 4.15 | 6.87 | 6.84 | 9.56 | 22.79 | 4.25 |
|  | 52-yr | 4.45 | 4.49 | 7.4 | 7.36 | 10.06 | 22.95 | 4.74 |
|  | 28-yr | 1.74 | 1.8 | 5.6 | 4.94 | 8.54 | 20.99 | 2.21 |
| Human (hypertension) | Normal | 5.78 | 5.84 | 8.63 | 8.61 | 11.22 | 24.2 | 6.07 |
|  | Mild | 3.47 | 3.5 | 6.21 | 6.2 | 9.53 | 21.85 | 3.64 |
|  | Sever | 6.02 | 6.03 | 8.58 | 8.59 | 11.16 | 24.02 | 6 |
| Animal (healthy) | Pig | 3.83 | 3.9 | 7.68 | 6.97 | 9.9 | 18.44 | 4.6 |
|  | Dog | -1.86 | -1.84 | 0.59 | 0.45 | 2.87 | 14.01 | -1.67 |

**TABLE III:** *AIC* calculated based on the developed fractional-order representation and standard ones for each subject.

|  |  | Model (A) | Model (B) | Model (C) | Model (D) | Model (F) | Model (E) | Model (G) |
| --- | --- | --- | --- | --- | --- | --- | --- | --- |
| Human (aging) | 68-yr | 3.01 | 3.02 | 5.18 | 5.14 | 7.3 | 17.7 | 3.12 |
|  | 52-yr | 3.32 | 3.36 | 5.7 | 5.66 | 7.8 | 17.87 | 3.61 |
|  | 28-yr | 0.61 | 0.67 | 3.9 | 3.25 | 6.28 | 15.9 | 1.08 |
| Human (hypertension) | Normal | 4.65 | 4.71 | 6.94 | 6.92 | 8.96 | 19.12 | 4.94 |
|  | Mild | 2.34 | 2.37 | 4.51 | 4.51 | 7.27 | 16.76 | 2.51 |
|  | Sever | 4.89 | 4.9 | 6.89 | 6.9 | 8.9 | 18.94 | 4.87 |
| Animal (healthy) | Pig | 3.67 | 3.74 | 7.44 | 6.73 | 9.58 | 17.72 | 4.44 |
|  | Dog | -2.02 | -2 | 0.35 | 0.21 | 2.55 | 13.3 | -1.83 |

**TABLE IV:** *AIC_C_* calculated based on the developed fractional-order representation and standard ones for each subject.

|  |  | Model (A) | Model (B) | Model (C) | Model (D) | Model (F) | Model (E) | Model (G) |
| --- | --- | --- | --- | --- | --- | --- | --- | --- |
| Human (aging) | 68-yr | 4.34 | 4.35 | 8.18 | 8.14 | 13.02 | 107.7 | 4.45 |
|  | 52-yr | 4.65 | 4.7 | 8.7 | 8.66 | 13.52 | 107.87 | 4.95 |
|  | 28-yr | 1.95 | 2 | 6.9 | 6.25 | 12 | 105.9 | 2.41 |
| Human (hypertension) | Normal | 5.99 | 6.04 | 9.94 | 9.92 | 14.68 | 109.12 | 6.27 |
|  | Mild | 3.68 | 3.7 | 7.51 | 7.51 | 12.98 | 106.76 | 3.84 |
|  | Sever | 6.22 | 6.23 | 9.89 | 9.9 | 14.62 | 108.94 | 6.21 |
| Animal (healthy) | Pig | 6.67 | 6.74 | 15.44 | 14.73 | 29.58 | -42.28 | 7.44 |
|  | Dog | 0.98 | 1 | 8.35 | 8.21 | 22.55 | -46.7 | 1.17 |

TABLE V presents the estimates of the unknown parameters of each fractional-order representation for each human and animal subject. Using *Model (A)* and *Model (B)*, for all the subjects, the fractional-order, *α*, is less than 1. These results demonstrate the fractional-order behavior within the apparent compliance. As in the estimation process, the estimate of *α* was only constrained to be positive, larger than zero (the lower bound is set to be 0 or the upper bound left unconstrained, equal to infinity). Therefore, this effect intends that the vascular system presents a viscoelastic behavior, not a purely elastic one. Actually, the fact that *α* ╪ 1 means that the fractional-order component involves both resistance and capacitance parts, as demonstrated mathematically in Eq. (9). The contributions from both the resistive and capacitive parts within the fractional-order capacitor are controlled through the fractional-order, *α*, allowing a profound physiological characterization. As the *α* moves to 1, the capacitance component becomes predominant and, hence the vascular mechanism functions as a pure elastic system, and as *α* goes to 0, the resistive portion expands, and so the vascular mechanism operates as a pure viscous system. By examining the estimates of the fractional orders of *Models (C)*, *(D)* and *(E)*, it is noticeable that for all the subjects, 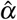 is higher than 1. Functionally, as *α* beats 1, the real part of the fractional-order capacitor impedance, *Z_r_*, converts negative, and hence it has the characteristic of a negative resistor producing power. Having a negative resistance in these models appears as compensation for the added static resistance in those representations. It is worth mentioning that the interest of constant resistor and/or capacitor in these fractional-order models is to account for the static viscosity and/or elasticity, respectively, while the fractional-order capacitor represents the ability of the arterial vessel to store blood dynamically.

**TABLE V:**
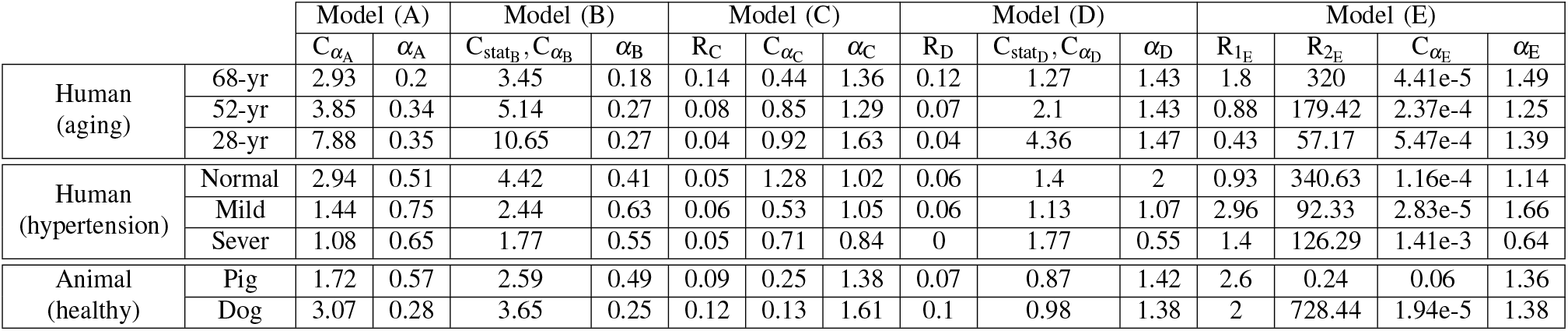
Parameter estimates of the fractional-order models for each subject.

### C. Convergence of the parameters

As shown in the parameter calibration algorithm 1, the estimate of *Θ^Model^* is 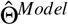 were found via the solution of the inverse problem of the estimated apparent compliance 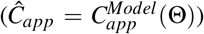 and the real one (*C_app_*). Initialized by *Θ*_0_ and using a nonlinear programming solver (*fmincon*), the inverse algorithm iteratively predicts the set of parameters 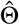 which minimizes the objective function, the root mean square error between the complex *C*_*app*_[*i*]__ and the model predicted *Ĉ*_*app*_[*i*]__(*Θ*) evaluated at the *i^th^* harmonic 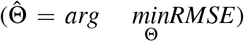. In this process, we constrained all the parameters to be positive to guarantee physical properties (Lower_*bounds*_ = [0], Upper*_bouds_* = [∞] and . Once a tolerance of error was reached, the convergence of the method is confirmed, the *fmincon* function exits and yields an output of the optimal set of model parameters estimates 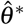. In order to study the convergence of the parameter estimations, 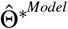, of the proposed fractional-order models, the estimation problem has been solved for each fractional-order model using different initial conditions.

Fig. A.1, in the appendix, shows the time history of the estimated values of the models’ parameters {Model (A); Model (B); Model (C); Model (D); Model (E);} during the optimization process starting from ten different initial conditions. In addition, Fig. A.2, in the appendix, displays the time history of the objective function (RMSE) during the optimization process starting from different initial conditions. Each color in these figures represents an iteration chain associated with a given initial condition configuration. Overall, it is clear that for each initial parameter configuration, after 100 iterations, the parameters and the *RMSE* converge to the same value. Conclusively, we can guarantee a good convergence of both the *RMSE* and parameters in the proposed fractional-order model’s calibration.

### D. Physiological consistency of the fractional differentiation order parameter

The fractional-order paradigm affords a concise alternative to characterize and quantify the biomechanical behavior of membranes, cells, and tissues. In fact, many studies have found that the fractional-order framework is particularly relevant in the area of biorheology characterization. This because many tissue-like materials present power-law responses to applied stress or strain. The power-law response has also been observed within the viscoelastic characterization of the aorta. *In-vivo* and *in-vitro* experiments and analysis showed the convenience of using fractional order viscoelastic model rather than the integer-order ones [21]. As shown previously, fractional differential order provides extra flexibility to model the apparent arterial compliance. It appears that the changes in the composition of the viscoelasticity of the whole vascular system are conveniently described in the fractional order of the model system. Based on the model formulation, the fractional parameter is convenient to describe the transition between viscosity and elasticity levels. Although the lack of enough real data, in this part, we investigate the interpretability and the physiological consistency of the fractional differentiation order.

For the ease of visualization, we present in Fig. 10, Fig. 12 and Fig. 11 the fractional differentiation order estimates, *α*, of each fractional-order model for the human and animal subject, respectively. In the case of human-hypertension, as shown in Fig. 10 we notice that the *α* values decreases from *Mild-hypertension* to *Severe-hypertension* conditions for all the proposed fractional-order models. These results are consistent with clinical analysis for hypertension and related risk factors. In fact, it is proven that stiff blood vessels and a lack of elasticity cause increased resistance to the flow of blood and high blood pressure. In addition, as shown in the previous sections, it is clear that as the fractional differentiation order decreases, the real part representing the resistive part increases and mirrors the stiffness behavior of the vessel. It is worth mentioning that in these results, we mark that *α* values in the normotensive case don’t show a good association with the other related-hypertension conditions based orders except for *Model(D)*. Conclusively, in the *human-hypertension* cases, the variation of the fractional-order is in compliance with the transition variation between viscosity and elasticity of the vascular vessel, which is revealed in the variation of the hypertension severity. Given the small number of observations, more data are needed to confirm this conclusion.

**Fig. 10:**
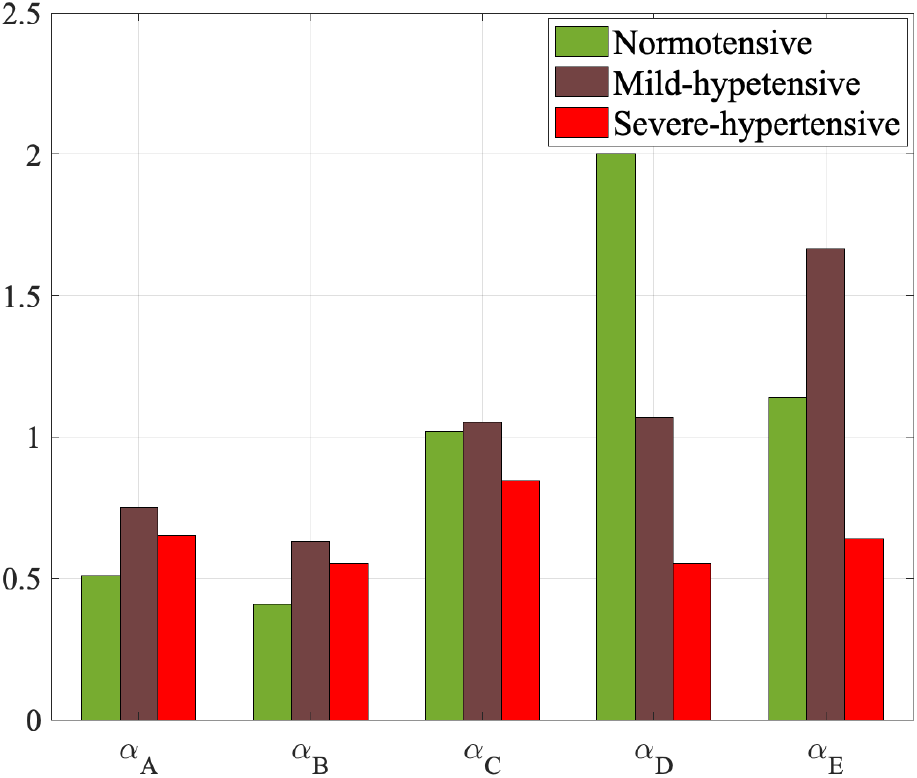
Estimated fractional differentiation order *α* using fractional-order models {Model (A), Model (B), Model (C), Model (D), Model (E)} for human hypertension.

**Fig. 11:**
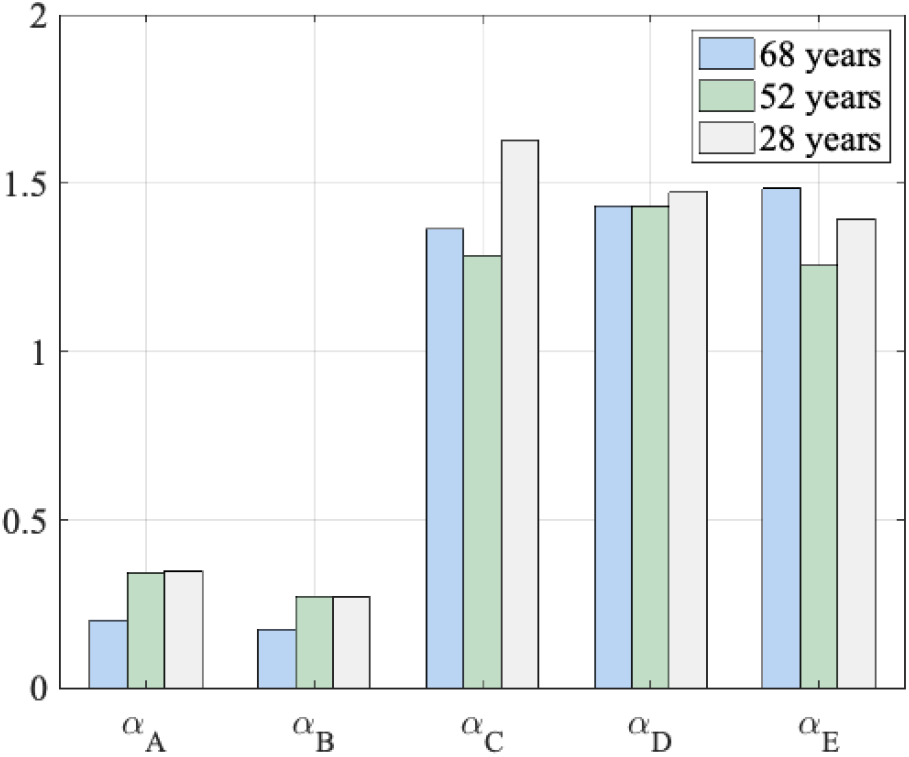
Estimated fractional differentiation order *α* using fractional-order models {Model (A), Model (B), Model (C), Model (D), Model (E)} for human-aging.

**Fig. 12:**
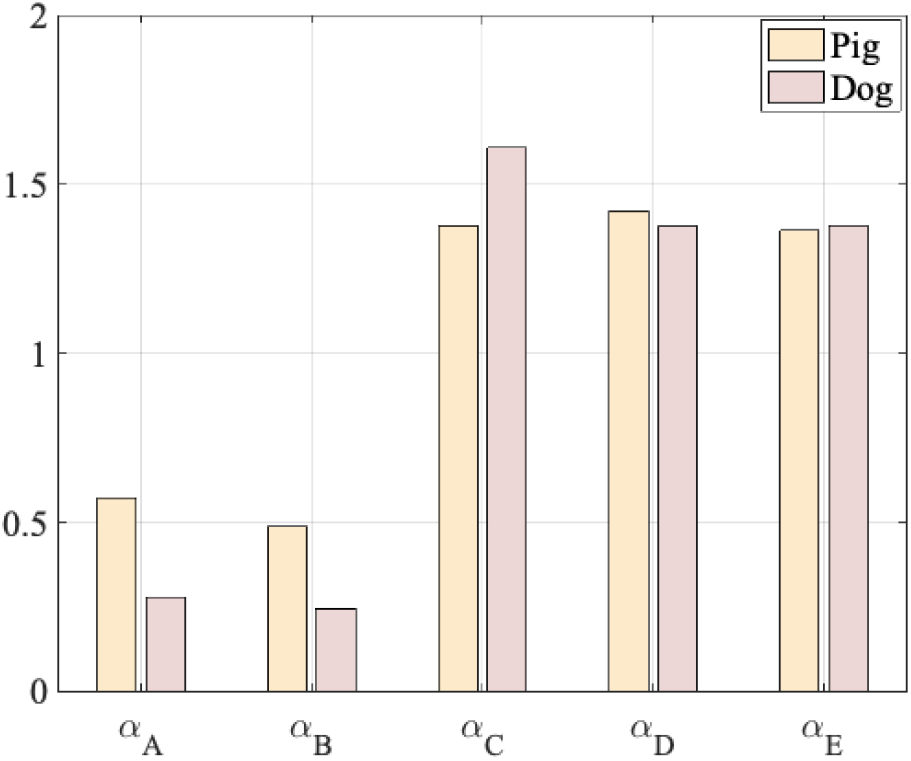
Estimated fractional differentiation order using fractional-order models {Model (A), Model (B), Model (C), Model (D), Model (E)} for animals (Pigs and Dogs) ones.

From Fig. 11, it is clear that *α_A_* and *α_B_* increase as the age decreases. This result is in coherence with what has been demonstrated in several human-aging studies. In fact, it is well recognized that the arterial vessel becomes stiffer with age. On the other hand, as we explained before, when *α* goes to 0, the resistive part increases within the fractional-order element, and the system behaves as a viscous element. For the other models based {*α_C_*, *α_D_*, and *α_E_* } where their values exceed 1, we can notice that from 68 years old subject to 52 years old one, the values of *α* decrease; however, the 28 years old subject presents the highest value. In this case, more real data are needed to affirm such conclusion and correlation between the evolution of the fractional differentiation order and age.

In Fig. 12, *α_A_* and *α_B_* of the dog is less than the ones of the pig. By checking the blood pressure waveform of these two animals in Fig. 3, it is clear that the pig’s systolic and diastolic blood pressure values are larger than the dog’s ones. Accordingly, the results can be interpreted as the increase of the hemodynamic values might be a consequence of increased vessel stiffness, leading to a decrease in *α*, for {*α_C_*, which exceeds 1, we notice that *α_C_* of the pig is less than the dog’s one, which is consistent with the previous results. However, { *α_D_* and *α_E_* } are approximately equal for both animals. The discussed results show the inherent benefits of using fractional-order elements in describing and characterizing the apparent arterial compliance. Fractional-order modeling offers an acceptable accuracy with a minimum number of parameters.

The analysis of the variation of the fractional differentiation order in *human-hypertension*, *human-aging* and animals points out the potential of this parameter to be adopted as a surrogate measure of the arterial stiffness or marker of cardiovascular diseases. Future clinical and experimental validations are required to prove the concept within a wide spectrum of normal and pathological cardiovascular conditions.

## VI. Example of arterial Windkessel model using fractional-order compliance

In order to evaluate the effect of integration the proposed fractional-order compliance representations within a complete arterial lumped parameter model, in this section we present a novel fractional-order Modified Windkessel model based on fractional-order capacitor. We validate the output pressure waveforms through forward fractional-order framework.

### A. Fractional-order modified Windkessel Model

The modified Windkessel model (MWK) is one of the simplest arterial representations that lumps the arterial network into two main compartments, proximal and distal, [39]. Taking into account that the proximal arteries close to the heart have different properties in comparison to the distal ones, MWK splits the total arterial compliance used in the original arterial Windkessel into two capacitances: proximal capacitor represents the compliance of the large arteries, which are commonly elastic and distal capacitor depicts the compliance of muscular arteries that are stiffer. Clinical studies demonstrated that distal compliance is very sensitive to vasodilatory experiments, a property apparent in distal arteries. other investigations have also shown that proximal compliance is reduced with aging and hypertension. The latest properties make these capacitances potential indicators of cardiovascular risk. Fig. 13 shows the circuit model of the fractional-order modified arterial Windkessel (F-MWK), which is similar to the MWK; however, instead of using integer-order ideal capacitors to represent the arterial compliances, simple fractional-order capacitors are employed. In this arterial lumped model, *C_α_* represents the compliance of large arteries close to the heart, while *C_β_* represents that of muscular arteries further away from the heart. *L* represents the inertance of the flowing blood. *R_p_* represents the peripheral resistance. *Q*(*t*) corresponds to the arterial blood flow and *P_ap_*(*t*) and *P_ad_*(*t*) denotes the proximal and distal pressure respectively.

**Fig. 13:**
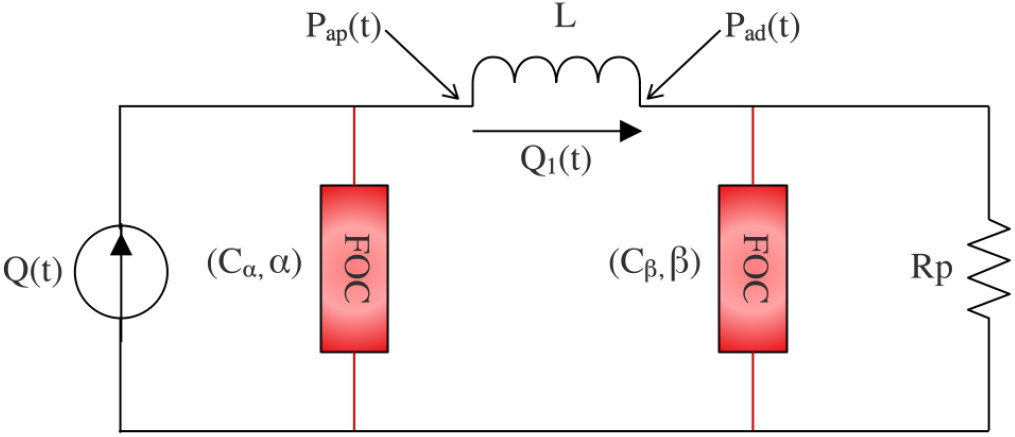
Fractional-order modified Windkessel circuit model. *C_α_* represents the compliance of large arteries close to the heart, while *C_β_* represents that of muscular arteries further away from the heart. *L* represents the inertance of the flowing blood. *R_p_* represents the peripheral resistance.

### B. F-MWK mathematical model

As the model comprises two fractional-order capacitors and an inductor, a state space representation with three states is written to describe the dynamic of the arterial system. Based on the Kirchhoff’s voltage and current laws, we obtain the following three equations:

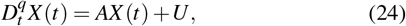

where,

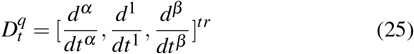

is the fractional-order derivative operator for all the states and

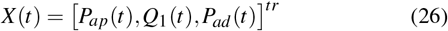

represents the state vector, (·)^*tr*^ denotes the transpose of the row vector.

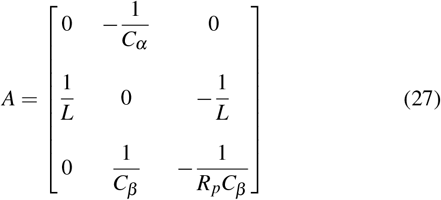

represent the parameters.

*U* is written as:

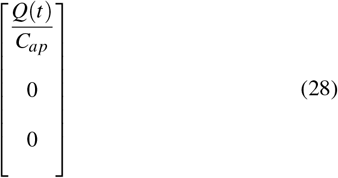

To implement the fractional-order derivative we used the following *Grünwald–Letnikov* (GL) formula [40]:

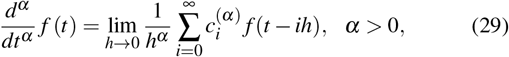

where *h* > 0 is the time step, 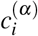 (*i* = 0, 1,…) are the binomial coefficients recursively computed using the following formula,

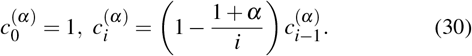

### C. Validation

The time validation of the proximal pressure waveforms *P_ap_* was performed through the proposed F-MWK using *in-vivo* human and animal database described in section IV. The optimizer algorithm uses the measured aortic root flow rate as an input and compute the required model parameters, {*C_α_*,*L*,*C_β_*,*R_p_*,*α*,*β*}, which minimize the pressure root mean square error (P-RMs), i.e. the difference between measured and calculated aortic root pressure as:

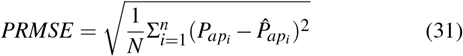

Where *N* denotes the number of samples per *P_ap_* pressure signal. To evaluate the performance of the estimation, we calculate the relative error, *R.E*.[%] defined as:

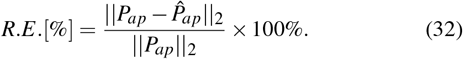

Fig. 14 shows the reconstructed blood pressure using the proposed F-MWK model along with the measured *in-vivo* blood waveform for each case of study, namely, human-aging, human-hypertension, and animal. In TABLE VI, the parameter estimates of the proposed F-MWK for each human and animal subject listen. We also present the *PRMSE* and *R.E*.[%] values for each subject. It is clear that the proposed model based on fractional-order compliance was able to capture the main features during the different phases of the cardiac cycle (systolic and diastolic phases) of the experimental data. The comparison between the obtained *R.E*. of the human aging and hypertension data indicates that the F-MWK reconstructs better the proximal pressure in the hypertension case. The best reconstruction is obtained in the case of the Pig blood pressure waveform with *R.E.* around 2.68%. In this case, the proposed model could catch all the features, including the dicrotic notch. The estimates of the fractional differentiation orders (*α*, *β*) and pseudo-capacitances (*C_α_*,*C_β_*) of the proximal and distal compliances, respectively, for all the human-aging, hypertension, and animal cases are presented in Fig. 15. These results show that as the age or the level of hypertension increases, the value of *α* and *β* decreases. We noticed the same behavior for the values of *C_α_* and *C_β_*. These results are in agreement with the clinical studies that prove that proximal compliance is reduced with aging. In fact, the decrease of the value of the fractional differentiation order and pseudo-capacitance implies an increase in the resistive part and a decrease in the compliance one.

**Fig. 14:**
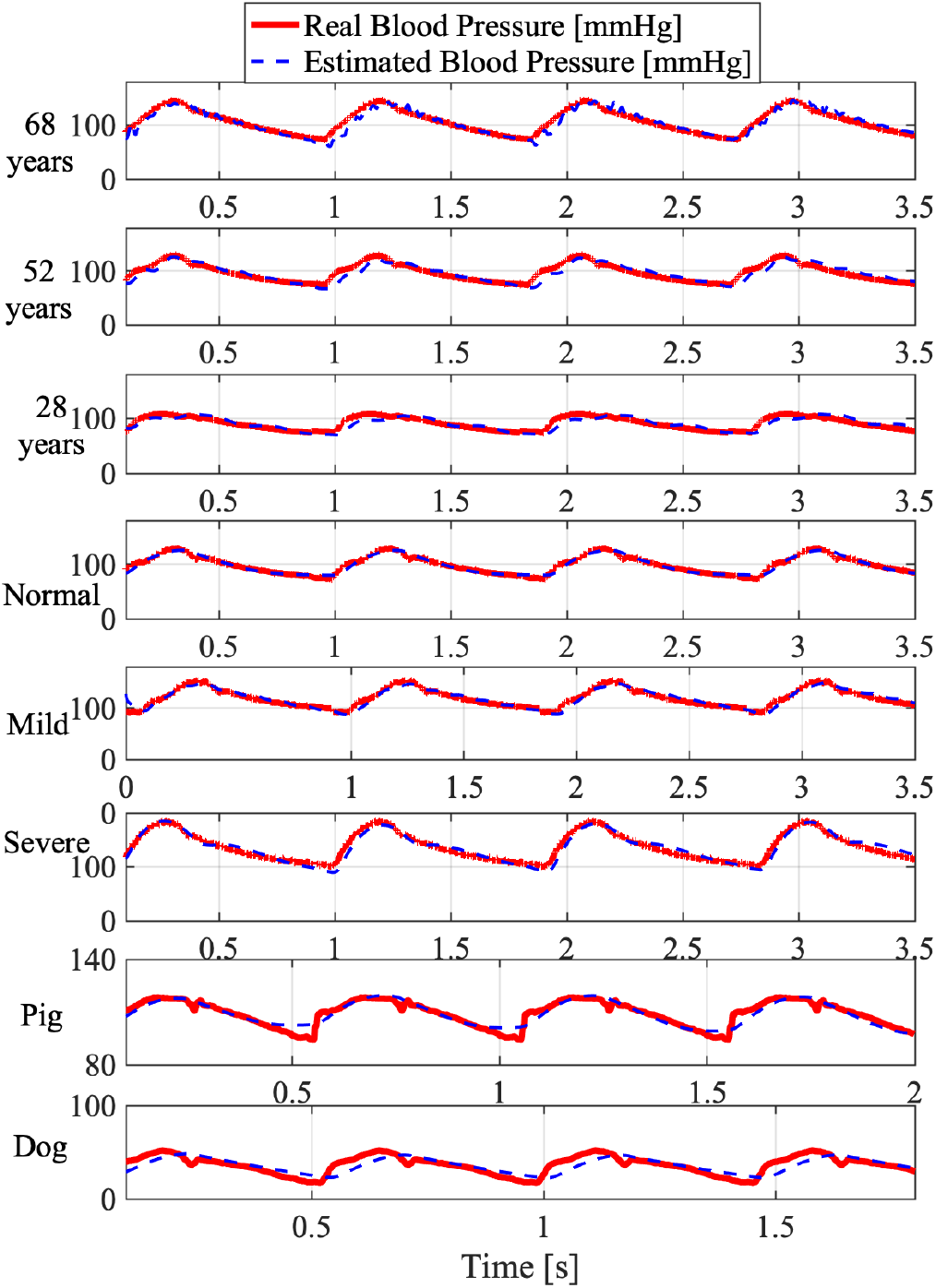
Estimated proximal blood pressure using the proposed fractional-order modified Windkessel model along with the experimental *in-vivo* human-aging and animal (Pigs and Dogs) ones.

**Fig. 15:**
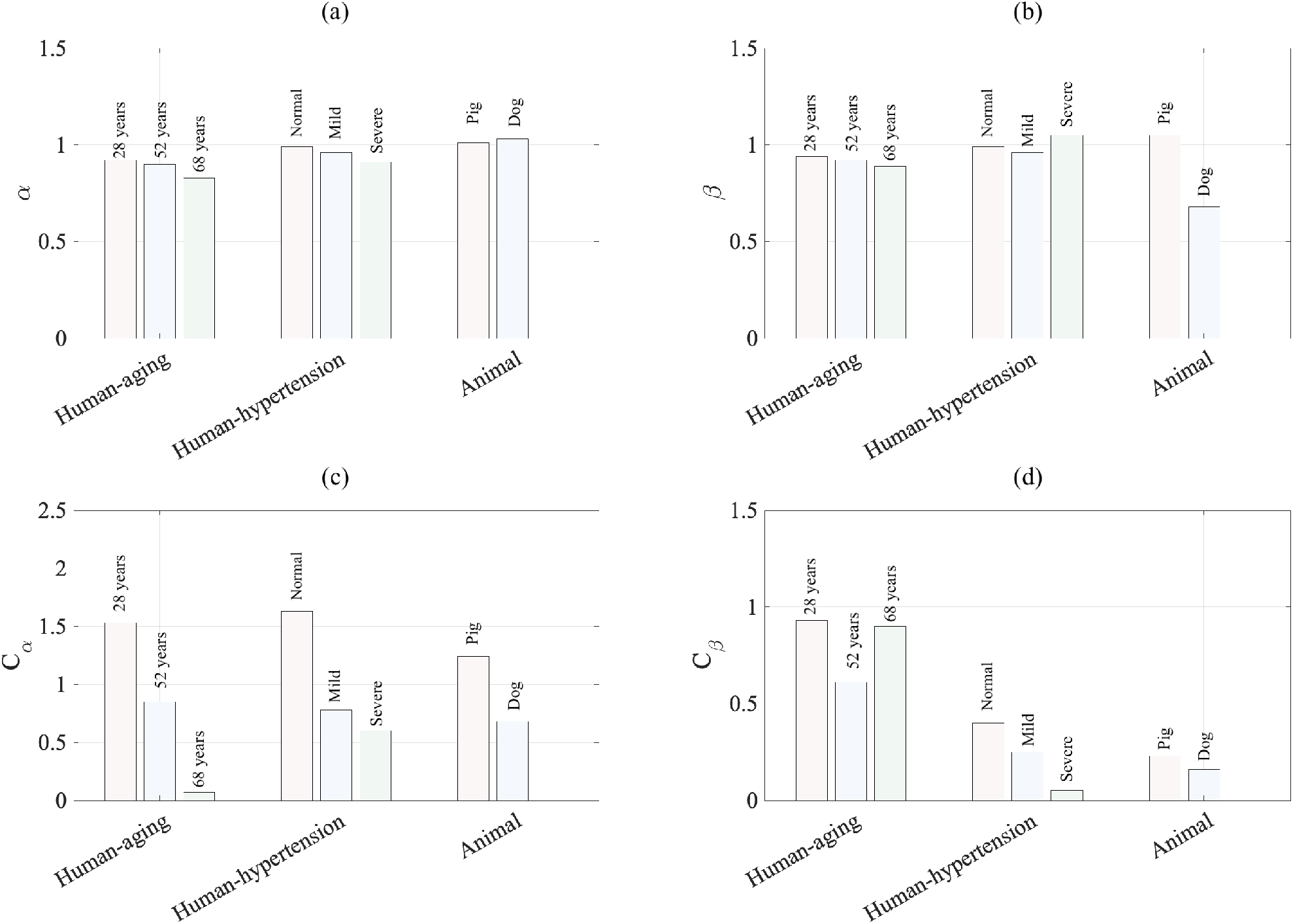
(a) Fractional differentiation order estimates, *α*, of the proximal compliance, (c) Fractional differentiation order estimates, *β*, of the distal compliance, (b) Pseudo-capacitance estimate of the proximal compliance *C_α_*, and (d) Pseudo-capacitance estimate of the distal compliance *C_β_*.

**TABLE VI:** Parameter estimates of the fractional-order modified Windkessel arterial models along with the *RMSEp* and the relative error *Re*[%] for each subject.

|  |  | Parameters estimates |  |  |  |  |  | Performance |  |
| --- | --- | --- | --- | --- | --- | --- | --- | --- | --- |
| | | $C_\alpha$ | $L$ | $C_\beta$ | $R_p$ | $\alpha$ | $\beta$ | $PRMSE$ | $R.E. [\%]$ |
| Human (aging) | 68-years | 0.07 | 0.0012 | 0.9 | 1.13 | 0.83 | 0.89 | 8.63 | 8 |
|  | 52-years | 0.85 | 0.0019 | 0.61 | 0.91 | 0.9 | 0.92 | 8.74 | 8.9 |
|  | 28-years | 1.53 | 0.0037 | 0.93 | 0.72 | 0.92 | 0.94 | 6.79 | 7.48 |
| Human (Hypertension) | Normal | 1.63 | 0.0592 | 0.4 | 0.88 | 0.99 | 0.99 | 4.01 | 4.05 |
|  | Mild | 0.78 | 0.0020 | 0.25 | 1.32 | 0.96 | 0.96 | 5.46 | 4.56 |
|  | Severe | 0.6 | 0.0853 | 0.05 | 1.96 | 0.91 | 1.05 | 7.8 | 5.65 |
| Animal (healthy) | Pig | 1.24 | 0.0342 | 0.23 | 1.98 | 1.01 | 1.05 | 2.92 | 2.68 |
|  | Dog | 0.68 | 0.0040 | 0.16 | 1.81 | 1.03 | 0.68 | 5.52 | 6.22 |

## VII. Conclusion

Arterial compliance is a vital determinant of the ventriculoarterial coupling dynamic. Its variation is detrimental to cardiovascular functions and is associated with heart diseases. Accordingly, assessment and measurement of arterial compliance are essential in diagnosing and treating chronic arterial insufficiency. Indices and surrogate measurements of arterial compliance present a non-invasive assessment of the vasculature’s health and can provide appropriate knowledge about an individual’s future risk of morbidity and mortality. The fractional-order behavior by mean of the power low response has been shown in the characterization of the collagenous tissues in the arterial bed, the arterial hemodynamic, the red blood cell membrane mechanics, and the heart valve cusp.

This paper investigates the fractional-order framework to characterize vascular compliance. Accordingly, we introduce five fractional-order lumped parametric representations to assess apparent arterial compliance. The proposed models vary in terms of the number of elements to characterize compliance’s dynamic. Every configuration contains a fractional-order capacitor (FOC) that accounts for the complex and frequency dependence characteristics of the compliance. FOC lumps both viscous and elastic properties of the vascular wall in one component, controlled through the fractional differentiation order (α) of FOC. To fully identify the proposed models, the unknown parameters and the fractional differentiation order were estimated using real hemodynamic data collected from human aging and hypertensive and animal (Pig and Dog) subjects. The developed parametric models produce an accurate reconstruction of the real data. In order to check the validity of the proposed concepts within a global arterial pattern, the simplest fractional-order compliance developed configuration consisting of a single FOC has been employed to account for the proximal and distal arterial compliance within a modified arterial Windkessel representation.

The validation results using real human and animal aortic blood pressure and flow data show a perfect reconstruction of the proximal blood pressure. In addition, the values of the compliances and theirs fractional differentiation orders were in agreement with the clinical results of the aging and hypertension implications. Conclusively, our investigation attests that the fractional-order modeling framework conveniently captures the dynamic capacity of the vascular system to store the blood. In addition, it shows that the fractional-order paradigm has a prominent potential to afford a novel alternative in assessing arterial stiffness.

In future work, we will propose a fractional-order distributed arterial network model. Besides, we will examine the effects of certain cardiovascular pathologies upon changes in the dynamic arterial compliance described by the fractional-order capacitor. To allow future authors to produce reproducible and easily-compared evaluations of the proposed approach, we provide a publicly available Matlab code used for the presented results and models calibration algorithm. The pre-processed real data employed in this work is also available at https://github.com/Bahloulm/Fractional-modeling-of-vascular-compliance.

## Acknowledgment

Research reported in this publication was supported by King Abdullah University of science and Technology (KAUST) with the Base Research Fund (BAS/1/1627-01-01).

## Appendix

### Convergence of the parameters

**Fig. A.1:**
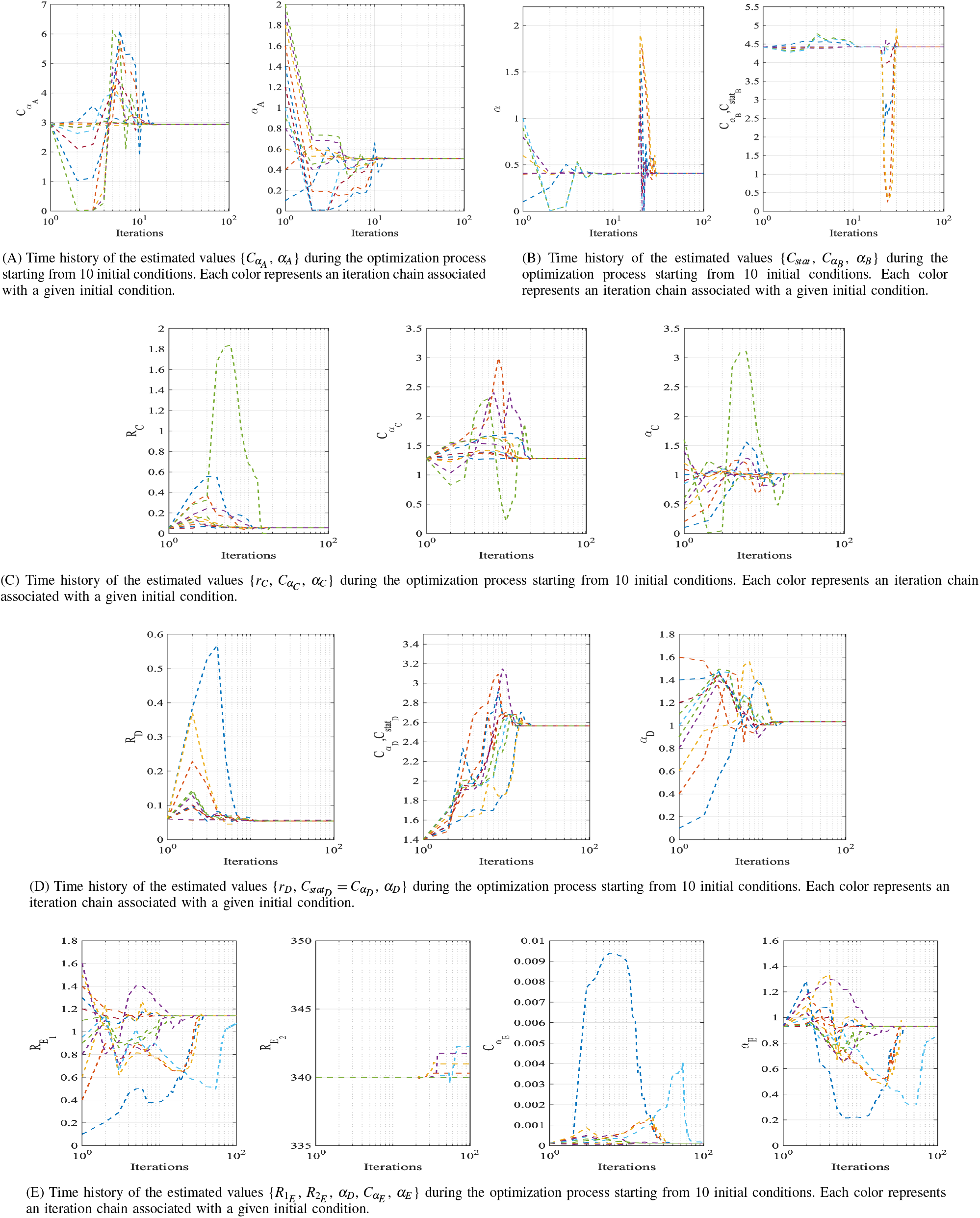
Time history of the estimated values of the models’ parameters, during the optimization process starting from 10 initial conditions. Each color represents an iteration chain associated with a given initial condition.

**Fig. A.2:**
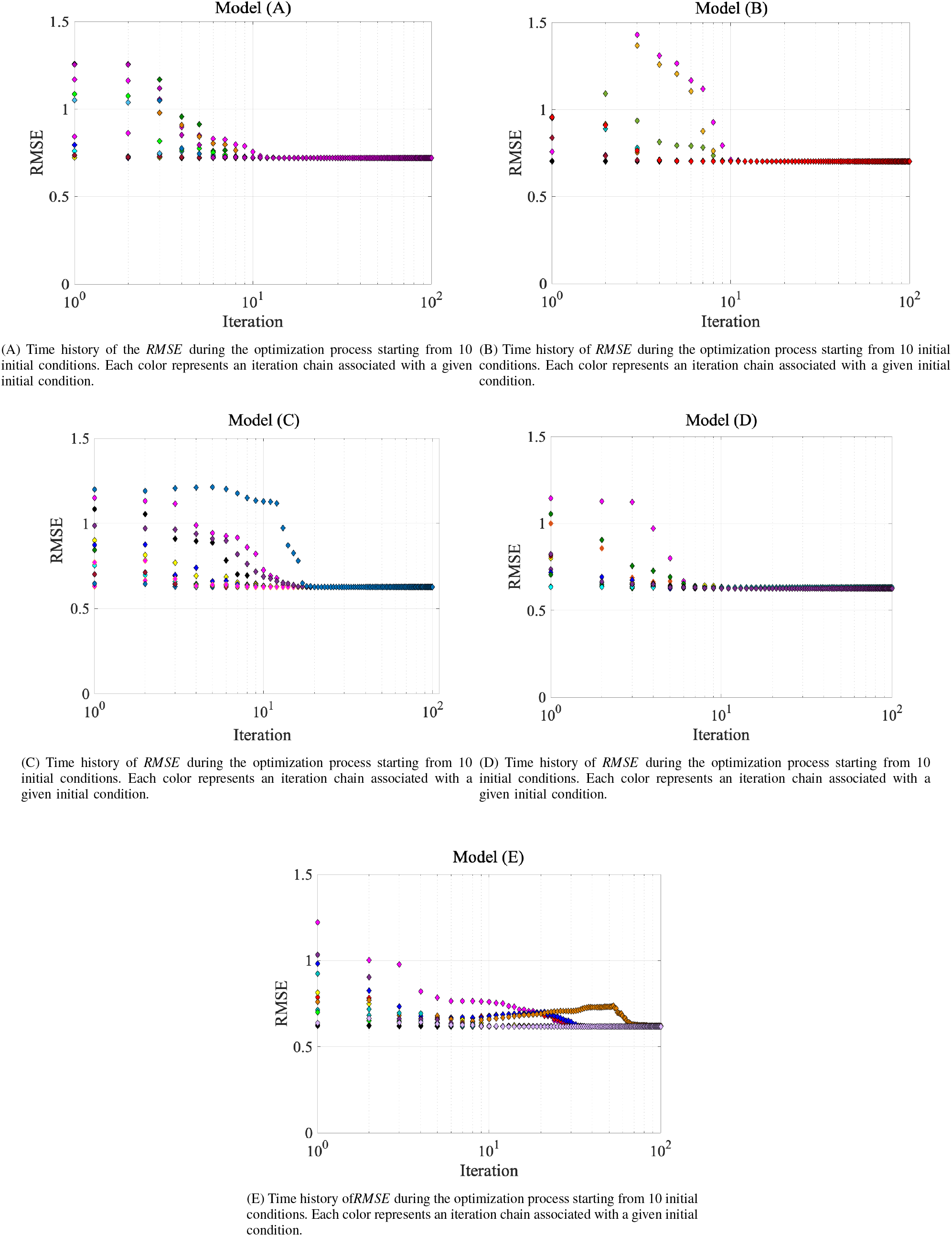
Time history of *RMSE*, during the optimization process starting from 10 initial conditions. Each color represents an iteration chain associated with a given initial condition.

